# Integrative cardiovascular dose-response to graded lower body negative pressure

**DOI:** 10.1101/2024.11.16.623934

**Authors:** Richard S. Whittle, Nathan Keller, Eric A. Hall, Safiyya Patanam, Bonnie J. Dunbar, Ana Diaz-Artiles

## Abstract

**Background:** Lower body negative pressure (LBNP) has been posited as a potential spaceflight countermeasure to counteract the physiological deconditioning related to fluid shifts in micro-gravity. However, open questions remain as to the magnitude of LBNP that should be applied. We systematically characterized the cardiovascular effects of LBNP and quantified the effect size of varied LBNP doses across different parts of the cardiovascular system.

**Methods:** Twenty-four subjects (12M, 12F) were exposed to graded LBNP from 0 mmHg to −50 mmHg in 10 mmHg increments, both in supine (0°) and 15° head-down tilt postures. We measured the steady-state response in a large range of variables, including those related to the systemic circulation, cardiovascular control, and hemodynamics of the eyes and neck.

**Results:** Building on the experimental data, dose-response curves were constructed using a Bayesian multivariate hierarchical modeling framework to quantify the effect size of every variable considered when subjected to LBNP. The methodology allows direct comparison of the variables and the underlying structural relationships between them. Further, we demonstrated the potential for LBNP to reduce jugular venous flow stagnation, which is considered one of the major health risks during human spaceflight.

**Conclusions:** The Dose-response curves and effect sizes generated from this research effort establish the most comprehensive framework available to date that characterizes physiological responses to LBNP. These results directly inform the development of countermeasures to mitigate the negative effects of spaceflight, including cardiovascular deconditioning, spaceflight-associated neuro-ocular syndrome, and venous thromboembolism events.

**Key Points Summary:**

- Lower body negative pressure (LBNP) has been posited as a potential countermeasure for multiple risks associated with spaceflight, including cardiovascular deconditioning, spaceflight-associated neuro-ocular syndrome, and venous thromboembolism events. However, the specific LBNP dose needed to mitigate these risks is still unknown.
- We constructed the LBNP dose-response curves in supine and 15° head-down-tilt positions to quantify the acute response of a large variety of hemodynamics, autonomic, ocular, and neck variables across a range of LBNP levels.
- We also used a Bayesian multivariate modeling framework to identify significant relationships between variables and to compare their effect sizes under different levels of LBNP.
- In addition, results indicate that LBNP is a promising countermeasure to reduce jugular venous flow stagnation occurring during microgravity exposure.
- This study constitutes the most comprehensive analysis of cardiovascular hemodynamics, autonomic, and cephalad response to LBNP to date. This framework informs the development of countermeasures to mitigate the detrimental effects of spaceflight.

## 1 Introduction

Future long-duration exploration missions will require novel countermeasure protocols to mitigate the degrading effects of the microgravity environment. Specifically relevant to the cardiovascular system is the headward fluid shift that occurs when entering in microgravity conditions. In particular, three of the risks directly related to the cephalad fluid shift identified by the NASA Human Research Program^1^ are: (1) the risk of cardiovascular adaptations contributing to adverse mission performance and health outcomes^2^; (2) the risk of spaceflight associated neuro-ocular syndrome (SANS)^3–5;^ and (3) the concern of venous thromboembolism^6–12^. Lower body negative pressure (LBNP) has a long spaceflight heritage since the Skylab Program in the 1970s^13,14^, both for human physiology research as well as a countermeasure to prevent post-flight orthostatic intolerance. Currently, Russian cosmonauts on the International Space Station (ISS) use the Chibis-M suit, developed in 2012, as a countermeasure prior to landing^15^. The Chibis protocol developed by the Institute for Biomedical Problems of the Russian Academy of Sciences (IBMP) is short, consisting of 2 min at −25 mmHg followed by 3 min at −35 mmHg^16^. The primary effect of LBNP is to pool blood down towards the feet, reducing venous return and introducing central hypovolemia^17^. Thus, LBNP is also used terrestrially to study the effects of acute hypovolemia and hemorrhagic shock^18,19^. Although LBNP does not restore hydrostatic gradients or affect tissue weight, this footward fluid shift could counteract the microgravity-induced cephalad fluid shift. LBNP has been demonstrated to effectively reduce intraocular pressure^20^, intracranial pressure (ICP)^21^, and optic nerve sheath diameter^22^ in multiple ground-based studies. In addition to its use as a countermeasure to prevent orthostatic intolerance, LBNP has also been posited as a potential countermeasure to mitigate cardiovascular degradation, SANS, and venous throboembolism (VTE) events during long-duration spaceflight^23^. To effectively develop successful LBNP protocols, it is important to fully quantify the effects of different levels of negative pressure on different aspects of the cardiovascular system. This is not just limited to the systemic hemodynamics, but rather a complete and comprehensive understanding of the hemodynamic and autonomic response. Additionally, and particularly important for SANS and VTE, it is also necessary to quantify the specific effects on the hemodynamics of the ocular system as well as the head and neck. Finally, there is a large amount of individual variation between crewmembers in terms of both anthropometry^24^ and LBNP tolerance^25^. Thus, it is further important to characterize any relationships between cardiovascular variables and easily measurable subject characteristics such as age, height, and weight. As one example, Buckey *et al.* have previously identified an association between IOP changes in microgravity and body weight^26^. This is particularly important as the profile of spaceflight participants broadens with the rise of commercial spaceflight^27^.

LBNP has been extensively studied in literature^15,17,18,21,22,25,28–36^. Multiple studies have previously looked at the difference between males and females, with the majority of them noting a difference in orthostatic tolerance^29,37–42^. For example, Patterson *et al.* highlight the importance of sex as a factor in the cardiovascular (CV) response to LBNP^41^. Fong *et al.* posited that the lower orthostatic tolerance observed in females (during centrifugation) were likely due to the combination of anthropometric factors, the vasoactive effects of sex hormones, and structural differences in cardiac anatomy^43^. However, none of the studies examined have considered the same range of hemodynamic, autonomic, and head/neck measurements considered in the present study. Further, there are some studies that examine tilt and LBNP as two separate interventions (e.g., Patterson *et al.*^41^, Greenwald *et al.*^20^, Ogoh *et al.*^44^). However, we could only find a few studies where both tilt and LBNP are considered together, i.e., LBNP during head-up tilt (HUT) or head-down tilt (HDT). In most of these cases, only a single value or small range of LBNP are examined^45–47^. Finally, the Bayesian workflow that we introduce in this study is noteworthy, with no previous studies examining the network of associations between variables.

The aim of this study is to construct dose-response curves to quantify the acute response of the cardiovascular system across a range of LBNP levels. We aim to encompass a wide range of systemic and autonomic parameters, as well as variables related to the head and neck, in order to provide a holistic picture of the integrated response to LBNP, particularly as it relates to potential use cases as a spaceflight countermeasure. Whilst other studies have considered the acute response to LBNP across multiple cardiovascular variables, we intend to focus on the spaceflight application and answer the question: “how much LBNP is required to compensate for the changes induced by a cephalad fluid shift?” in any given variable of interest. We further intend to quantify any sex-dependent differences in LBNP response. Finally, in the process of analyzing our data, we developed a novel workflow for the construction of dose-response curves using Bayesian multivariate analysis. This multivariate framework captures the relationships between all of the measured variables, as well as subject characteristics such as age, height, and weight. Such a methodology could be expanded beyond the cardiovascular system to encompass other organ systems. Together, these results lead to a greater understanding of LBNP as a potential spaceflight countermeasure, informing the development of future protocols.

## 2 Methods

### 2.1 Subjects and Study Approval

Twenty-four healthy, recreationally active subjects (12 male, 12 female) between 22 and 42 years old were recruited from the Texas A&M University System to participate in the study. Subjects were matched for age and body mass index (BMI) between the male and female groups. Sample size and the number of pressure levels required were determined based on a power curve analysis of pilot data. Subject characteristics (mean ± SD) are presented in Table 1. Table 1 also gives the Bayes Factor (*BF*_10_), which is a Bayesian term that represents the strength of the evidence supporting differences between the male and female groups. Thus, the larger the *BF*_10_, the more evidence to reject the null hypothesis that the standardized effect size of the difference in a characteristic between groups (*d*) is negligible (where negligible is defined by a region of practical equivalence, ROPE, of ±0.1)i, such that:

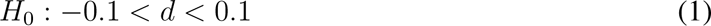

**Table 1:**
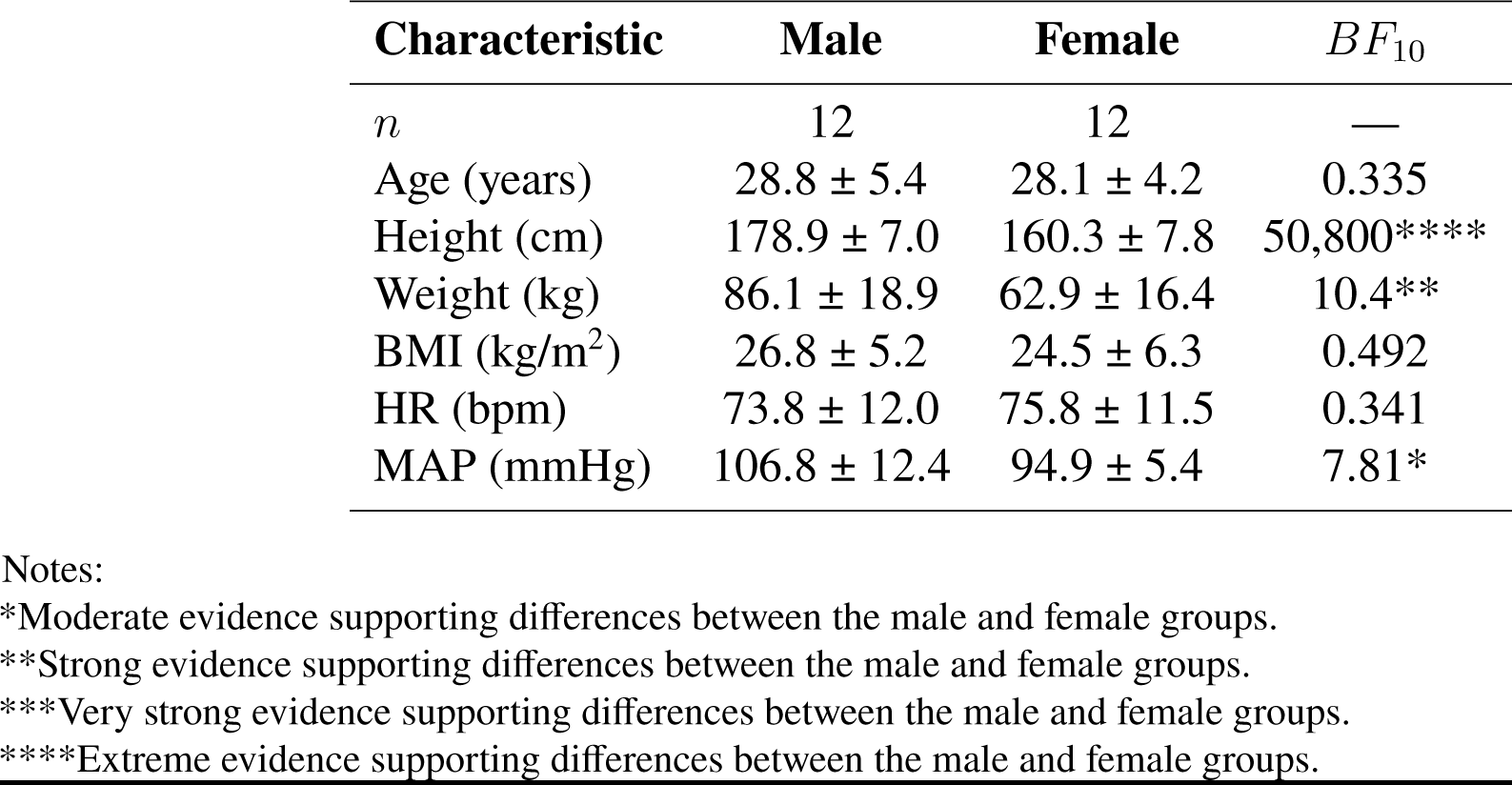
Characteristics of the 24 recreationally active subjects (12 male, 12 female) who participated in the study. Characteristics were recorded during baseline session prior to testing sessions. Data are reported as mean ± SD where appropriate. *BF*_10_ represents the strength of the evidence supporting differences between the male and female groups. Abbreviations: BMI, body mass index; HR, heart rate; MAP, mean arterial pressure.

| Characteristic | Male | Female | $BF_{10}$ |
| --- | --- | --- | --- |
| $n$ | 12 | 12 | — |
| Age (years) | $28.8 \pm 5.4$ | $28.1 \pm 4.2$ | 0.335 |
| Height (cm) | $178.9 \pm 7.0$ | $160.3 \pm 7.8$ | 50,800**** |
| Weight (kg) | $86.1 \pm 18.9$ | $62.9 \pm 16.4$ | 10.4** |
| BMI ($\text{kg}/\text{m}^2$) | $26.8 \pm 5.2$ | $24.5 \pm 6.3$ | 0.492 |
| HR (bpm) | $73.8 \pm 12.0$ | $75.8 \pm 11.5$ | 0.341 |
| MAP (mmHg) | $106.8 \pm 12.4$ | $94.9 \pm 5.4$ | 7.81* |

Notes:
\*Moderate evidence supporting differences between the male and female groups.
\*\*Strong evidence supporting differences between the male and female groups.
\*\*\*Very strong evidence supporting differences between the male and female groups.
\*\*\*\*Extreme evidence supporting differences between the male and female groups.

Prior to participating in the study, subjects completed a questionnaire designed to identify any exclusion criteria, including current use of any cardiac, blood pressure, muscle relaxant, anticoagulant, or stimulant medications, thyroid disease, chronic cardiovascular pathologies, extreme obesity, history of hypertension, or possible pregnancy. Each subject received written and verbal explanations of the study protocols and gave written informed consent to participate in the experiment. All procedures performed in the study were in accordance with the 1964 Helsinki Declaration and its later amendments. The study protocol was approved by the Texas A&M Human Research Protection Program with Institutional Review Board number IRB2020-0724F.

### 2.2 Experimental Design and Testing Protocol

The experiment was designed as a counterbalanced, within-subjects, experiment such that every subject experienced every pressure level and posture. Subjects were exposed to graded LBNP from 0 mmHg to −50 mmHg in 10 mmHg increments (in a progressive order) in two separate postures: 0° supine (face-up) and 15° head-down tilt (HDT) supine. The procedure was identical for each posture. Experimental sessions took place on two separate days within a two-week period. In each of the two experimental sessions, subjects were tested either in the 0° supine position or in the 15° HDT position (order counterbalanced). Additionally, in the first session, baseline measurements were collected in a seated posture prior to the main testing. To control for potential circadian effects, all sessions were scheduled in the morning at approximately the same time. In addition, subjects were asked to refrain from drinking caffeine and exercising prior to each test session.

In a single experimental session (0° supine or 15° HDT), subjects were placed in a lower body negative pressure chamber (Technavance, Austin, TX) initially at 0 mmHg differential with atmospheric pressure. Continuous measurements of blood pressure and electrocardiography were recorded throughout the test. Subjects initially remained at rest for a period of six-minutes to allow any hemodynamic transients to settle. After the rest period, an inert gas rebreathing device was used to collect discrete measurements of cardiac parameters. Following this, measurements of ocular tonometry, ultrasonography, and non-invasive measurement of internal jugular venous pressure were collected from the subjects. The total procedure at a single pressure level lasted for approximately 12 minutes. The LBNP pressure level was then increased (more negative) by 10 mmHg and the entire process repeated, starting with the six-minute resting period. The total protocol included six pressure levels: 0 mmHg, −10 mmHg, −20 mmHg, −30 mmHg, −40 mmHg, and −50 mmHg. The procedure for the seated baseline conducted on the first experimental session was identical to the procedure for a single pressure level.

Testing was discontinued immediately if subjects experienced discomfort, or presented physiological markers of presyncope such as unrestrained rising heart rate, falling blood pressure, and/or profuse perspiration. In the 0° supine position, presyncope was reached in seven subjects at −40 mmHg (*n* = 2, both female) and −50 mmHg (*n* = 5, 4 female, 1 male). In the 15° HDT position, presyncope was reached in two subjects (both female) at −40 mmHg (*n* = 1, one of them also reached presyncope at −40 mmHg whilst in 0° supine) and −50 mmHg (*n* = 1, also experienced presyncope at −50 mmHg whilst in 0° supine). After discontinued application of LBNP, no subjects experienced lasting symptoms. The remainder of the data for these subjects up to the point of discontinuation are included in the results. All other subjects completed the full protocol and experienced no adverse effects.

### 2.3 Dependent Variables

Dependent variables included 11 hemodynamic metrics, seven autonomic indices, and eight measures related to the head/neck/eyes. The hemodynamic measurements considered were: 1) heart rate (HR, bpm); 2) stroke volume (SV, ml); 3) cardiac output (CO, l/min); 4) oxygen consumption (VO2, l/min); 5) systolic blood pressure (SBP, mmHg); 6) diastolic blood pressure (DBP, mmHg); 7) rate pressure product (RPP, mmHg/min), used as a metric for myocardial stress and energy consumption^50^; 8) myocardial oxygen supply:demand index (MO, calculated as the ratio of the diastolic pressure time interval to the systolic pressure time interval, DPTI/SPTI, no units)^51^; and 9) total peripheral resistance (TPR, mmHg.s/ml). Additionally, two body weight normalized indices were collected: 10) stroke index (SI, ml/m^2^); and 11) cardiac index (CI, l/min/m^2^).

Following the recommendations of the Task Force of the European Society of Cardiology and the North American Society of Pacing and Electrophysiology^52^, we collected four time-domain autonomic indices and three frequency-domain autonomic indices. These measurements were: 1) the standard deviation of the NN intervals (SDNN, ms); 2) heart rate variability triangular index (HRVTi, no units); 3) the root mean square of direct differences of the NN interval (RMSDD); 4) baroreflex sensitivity (BRS, ms/mmHg); 5) normalized spectral power density in the low frequency (0.04–0.15 Hz) band (LFNorm, no units); 6) normalized spectral power density in the high frequency (0.15–0.4 Hz) band (HFNorm, no units); and 7) the ratio between low frequency and high frequency power spectral densities (LF/HF, no units). SDNN and HRVTi represent heart rate variability incorporating sympathetic and parasympathetic effects, RMSDD and HFNorm are closely correlated with parasympathetic activity^53^, LFNorm represents sympathetic activity, LF/HF represents sympathovagal balance, and BRS represents a measure of total autonomic control via the arterial baroreflex^52,54,55^.

In relation to the head and neck, the following eight measurements were collected (or calculated): 1) intraocular pressure (IOP, mmHg); 2) ocular perfusion pressure (OPP, mmHg); 3) internal jugular vein cross-sectional area (A_ĲV_, mm^2^); 4) internal jugular vein pressure (ĲVP, mmHg); 5) internal jugular vein flow pattern (ĲVF, grade − see discussion in Section 2.4 below); 6) common carotid artery cross-sectional area (A_CCA_, mm^2^); 7) common carotid artery peak systolic velocity (PSV, cm/s); and 8) common carotid artery end diastolic velocity (EDV, cm/s). Head and neck measurements were collected on both the right and left sides.

### 2.4 Instrumentation and Data Collection

Hemodynamic measurements were collected using two instruments: an Innocor inert gas rebreathing device (Cosmed: The Metabolic Company, Rome, Italy) and a Finapres NOVA (Finapres Medical Systems B.V., Enschede, the Netherlands). Full calibration was performed on devices daily, and ambient data calibrations were also performed prior to each subject test (mean ± SD: temperature 24.4 ± 1.2°C, relative humidity 46.7 ± 5.8%, pressure 753.6 ± 4.2 mmHg). The inert gas rebreathing method was used to obtain noninvasive measures of pulmonary blood flow by analyzing the changing concentrations of a soluble gas (nitrous oxide, 5%) and an insoluble gas (sulfur hexafluoride, 1%) in an oxygen-enriched air mixture over 5-6 breaths. The mixture is rebreathed using a bag for approximately 30 seconds. During the rebreathe, subjects inspired and expired at a rate of 20 breaths per minute, following this rhythm with a metronome (respiration at all other times was at a normal relaxed respiration rate). After each rebreathe, the gas concentration traces were visually inspected by a trained operator to ensure correct function of the device. Further details on the inert gas rebreathing methodology can be found in Whittle *et al.*^56^. Finapres data (finger arterial pulse contour waveform and 5-lead electrocardiogram) were collected continuously throughout the protocol with pressure corrected to heart level with a hydrostatic height sensor placed laterally on the mid-coronal plane at the fifth intercostal space. At each pressure level, the Finapres pressure waveform was calibrated with a discrete blood pressure measurement using a brachial sphygmomanometer. Autonomic indices were derived from the Finapres ECG trace and beat-to-beat RR.

Measurements of intraocular pressure (IOP) were obtained at each pressure level using a contact tonometer (IC200, iCare, Vantaa, Finland). Values presented are the mean of the central four of six measures (i.e., a trimmed mean) to account for arterial and respiratory fluctuation. Ocular perfusion pressure (OPP) was manually calculated for each subject from their IOP and MAP at eye level (OPP = MAP_eye_ - IOP). MAP_eye_ was calculated from MAP corrected for: a) the hydrostatic pressure difference between heart and eye-level, and b) the perpendicular distance from mid-coronal plane of the body to anterior placement of the globe of the eye, using a procedure previously described in Petersen *et al.*^57^ and Hall *et al.*^58^.

Internal jugular vein pressure, ĲVP, was obtained by manually compressing the ĲV (around the C3 vertebral level) with a VeinPress (Compremium, Bern, Switzerland) manometer attached to the head of an ultrasound probe (VScan Extend, GE Healthcare, Chicago, IL). The VeinPress device was zeroed prior to each measurement. Pressure was recorded at the point at which the walls of the ĲV vessel were just about to touch each other. When this occurred, the pressure reading was allowed to stabilize for two seconds to counter any inertial effects. Two ĲVP measurements were collected at each LBNP-position-side combination, and the final ĲVP in that condition was calculated as the average of the two measurements.

All other measurements of the carotid arteries and jugular veins were collected using a Butterfly iQ+ ultrasound device (Butterfly Network Inc., Burlington, MA). Four separate images were obtained from each side of the subject in each experimental condition (i.e., pressure-position combination). Two images captured a transverse view of the CCA and ĲV, respectively, collected approximately 30 mm inferior to the CCA bifurcation point (around the C3 vertebral level) at end-diastole. Two trained operators, acting independently, manually identified and circumscribed the CCA and ĲV on each image to calculate A_CCA_ and A_ĲV_ based on pixel count. If the two measured areas from the different operators differed by less than 10%, the final A_ĲV_ in that condition was calculated as the average of the two independently measured areas. However, if the measured area differed by more than 10%, a third operator repeated the circumscription and the final A_ĲV_ in that condition was calculated as the average of the three independently measured areas.

The final two images captured spectral pulse-wave Doppler flow of the CCA and ĲV, respectively. For the CCA flow, an envelope tracing algorithm was applied, modified from an algorithm developed by Wadehn and Heldt^59^. The output of this algorithm was used to calculate average peak systolic velocity (PSV) and end-diastolic velocity (EDV). ĲV was binned into categories representing four flow regimes as defined by Marshall-Goebel *et al.*^10^. These flow grades represent: Grade 1 − forward flow that never returns to zero; Grade 2 − forward flow that may return to zero; Grade 3 − stagnant flow characterized by equal forward and retrograde flow; and Grade 4 − predominantly retrograde flow. No instances of grade 4 flow were observed. Examples of ĲV blood flow velocity waveform grades 1, 2, and 3 found in the present study are presented in Figure 1.

**Figure 1:**
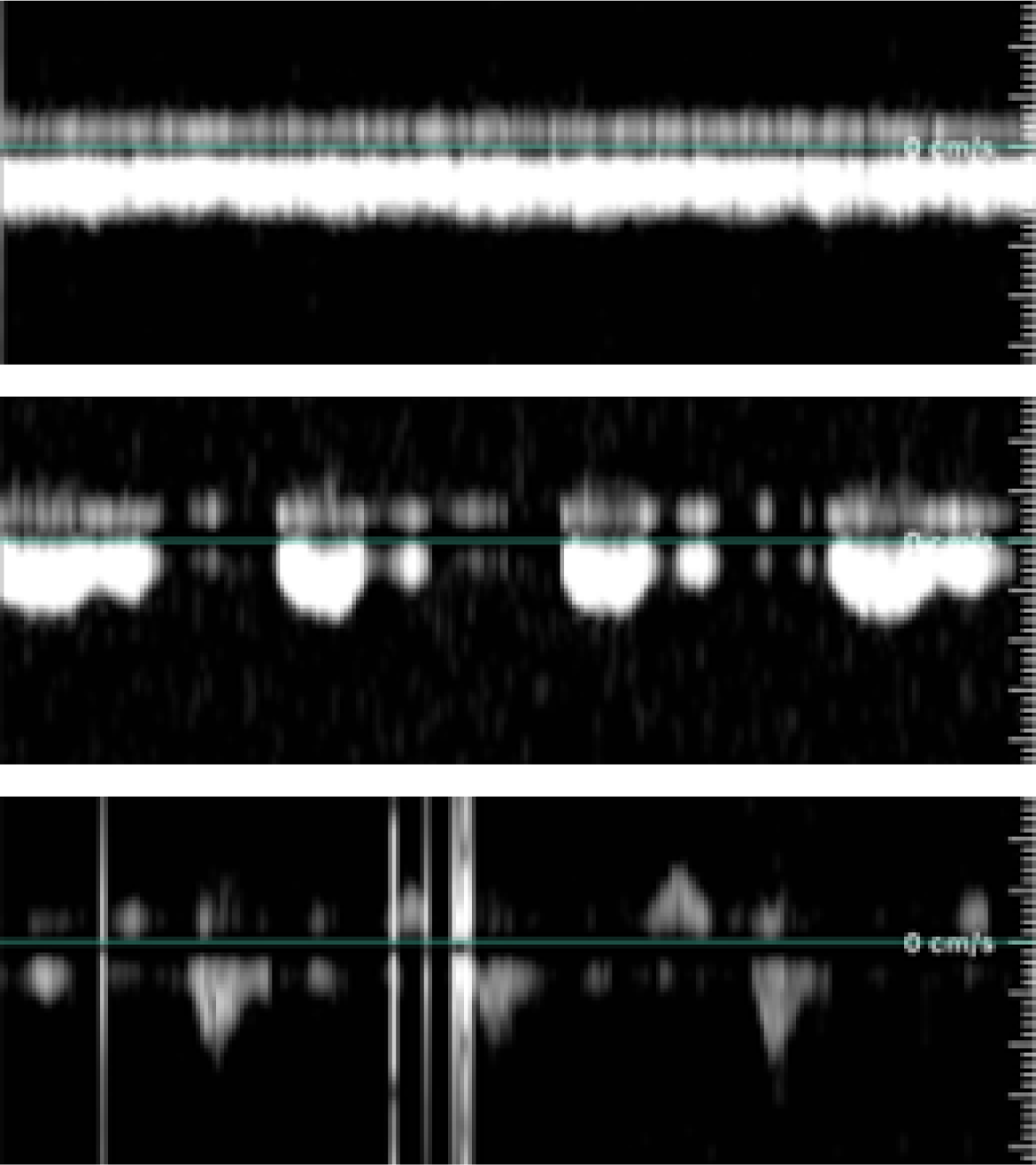
Examples of internal jugular vein blood flow velocity waveform grades found in the present study. (Top) Grade 1: forward flow that never returns to zero; (Middle) Grade 2: forward flow that may return to zero; (Bottom) Grade 3: stagnant flow characterized by equal forward and retrograde flow. Flow grade regimes are as defined by Marshall-Goebel *et al*.^10^.

### 2.5 Statistical Analysis

We generated dose-response curves using a Bayesian workflow in order to capture dependent structural relationships between all of the variables measured via a multivariate analysis. In the next few paragraphs, we fully and comprehensively report our methodology following the Bayesian Analysis Reporting Guidelines (BARG) given by Kruschke^60^.

All variables measured (with the exception of ĲVF, described separately below) exhibited an approximately linear response to graded LBNP. A single Bayesian robust multivariate, hierarchical regression model was used to estimate the effects of LBNP pressure, sex, and position (0° supine or 15° HDT) on all dependent variables. The multivariate model is presented in Equation 2:

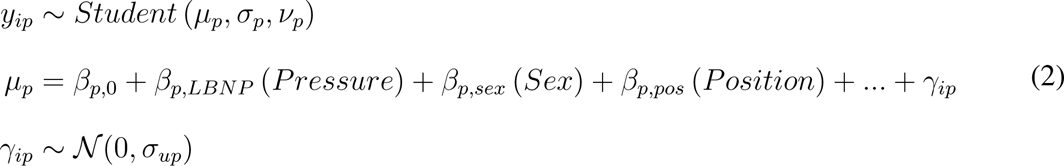

where, for each dependent variable *p* outlined in Section 2.3, *y* represents the standardized response, *σ* represents the population level standard deviation, *ν* represents the degrees of freedom, and *µ* represents the mean value. *µ* is further described by *β*_0_, *β_LBNP_*, *β_sex_*, and *β_pos_*, representing the coefficients for the independent variables, and *γ_i_* representing a group-level intercept for subject *i*, distributed normally with mean 0 and standard deviation *σ_u_* represents the inclusion of any additional terms due to interaction effects (see below). *Pressure* is an index variable representing the LBNP level, with 0 indicating 0 mmHg through to 5 indicating −50 mmHg, such that *β_LBNP_*gives the effect size of a decrease in 10 mmHg. *Sex* represents a contrast coded variable (−0.5 = Female, 0.5 = Male), such that *β_sex_* gives the effect size of the increase in *y_p_* in males over females. *Position* is a categorical variable with 0 representing 0° supine and 1 representing 15° HDT, such that *β_pos_* gives the effect size of a change from 0° supine to 15° HDT.

Prior to constructing the dose-response curves, all dependent variables (except ĲVF) were standardized to have a mean of 0 and a standard deviation of 0.5. This standardization was implemented for three principal reasons. First, the prior choice was greatly simplified, as priors on the same scale could be used for all dependent variables. Second, the computational efficiency of the Bayesian calculation was greatly improved. Finally, and most importantly, this process allowed for the comparison of the magnitude of the effect sizes across different variables. Results of our modeling efforts are presented both in this standardized form, and also back-transformed to the original scales of measurement. Finally, correlations were modeled to exist between the dependent variables, *p*, such that the covariance matrix of all the *σ_u_* for the group-level intercepts was described by a Cholesky decomposition.

For each dependent variable *p*, Bayes Factors analysis of a simple univariate regression was used to determine any additional interaction effects to be included in the model, with a Bayes factor of 3 (i.e., substantial evidence) used as the decision rule in favor of a more complicated model^61^.

For the eight variables related to the head and neck, an additional independent contrast coded variable, *Side* (and any appropriate interactions, determined using Bayes Factors), was included, such that *β_side_* gives the effect size of the increase in *y_p_* in the right side over the left side.

In the case of ĲVF and due to the nature of the data, we implemented an ordinal logistic regression model. For this model, the dependent variable was flow grade (from 1 to 3). The model used a binomial distribution with a logistic (logit) link, presented in Equation 3. *Pressure*, *Sex*, *Position*, and *Side* remained as the predictor variables and the group-level intercept was allowed to correlate with the remainder of the dependent variables as described above. In this case, the coefficients, *β*, of the independent variables represent the log odds of either a grade 2 or grade 3 flow pattern with respect to a grade 1 pattern. Thus, *e^β^* represents the odds ratio (OR). The ĲVF model is further described below:

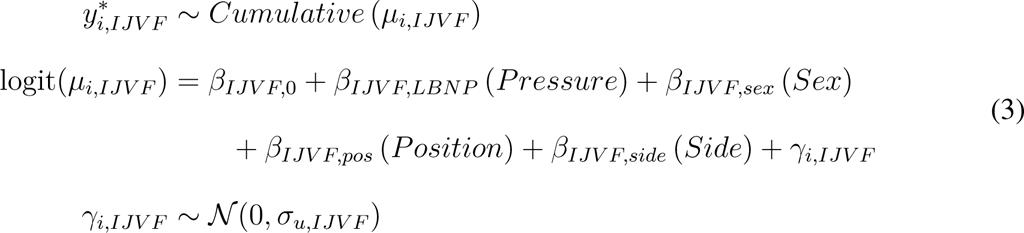

*where* 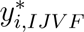 is the latent variable for the ĲV blood flow velocity waveform pattern for subject *i; µ_i,IJV F_* is the linear predictor (with a logit link function); *β_IJV F,_*_0_, *β_IJV F,LBNP_*, *β_IJV F,sex_*, *β_IJV F,pos_*, and *β_IJV F,side_* are the coefficients for the intercept, LBNP *Pressure*, *Sex* (male or female), *Position* (0° supine or 15° HDT), and *Side* (right or left), respectively; *γ_i,IJV F_* is the group-level intercept for subject *i*; and *σ_u,IJV F_* is the standard deviation of the group-level intercept. Due to significant heterogeneity (i.e., the variance of the data increasing with stronger LBNP), a distributional regression model was used such that *σ_IJV P_* was allowed to vary with the pressure level as indicated in Equation 4:

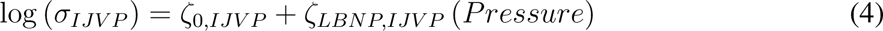

where *ζ*_0*,IJV*_ *_P_* and *ζ_LBNP,IJV P_* are the intercept and slope of the log of the *σ_IJV P_*, respectively.

Finally, in order to determine the multivariate relationship between the variables measured and the subject characteristics, standardized *Age*, *Height*, *W eight*, and *BMI*, were added to the multivariate regression model as dependent variables, defined as a unique group-varying intercept in the form of Equation 5:

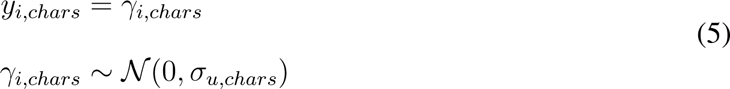

where *y_i,chars_*is the characteristic (*Age*, *Height*, *W eight*, or *BMI*) of subject *i*; and *γ_i,chars_* is a group-level intercept with standard deviation *σ_u,chars_*.

Table 2 presents the form of the regression model for all dependent variables, *p*.

**Table 2:** Bayesian multivariate regression model used to construct the dose-response curves for the cardiovascular response to LBNP. Distributions, main effects, and additional effects (interaction or distributional parameters) for each dependent variable are included in the table. All dependent variables are combined into a single matrix and analyzed using a single, large regression model as detailed in the text.

| Measure | Distribution | Main Effects | Additional Effects<sup>†</sup> |
| --- | --- | --- | --- |
| HR | *Student* | *Pressure, Sex, Position* | — |
| SV | *Student* | *Pressure, Sex, Position* | *Pressure × Sex* |
| CO | *Student* | *Pressure, Sex, Position* | — |
| VO2 | *Student* | *Pressure, Sex, Position* | *Sex × Position* |
| SBP | *Student* | *Pressure, Sex, Position* | — |
| DBP | *Student* | *Pressure, Sex, Position* | — |
| RPP<sup>‡</sup> | $\mathcal{N}$ | *Pressure, Sex, Position* | — |
| MO | *Student* | *Pressure, Sex, Position* | — |
| TPR | *Student* | *Pressure, Sex, Position* | — |
| SI | *Student* | *Pressure, Sex, Position* | — |
| CI | *Student* | *Pressure, Sex, Position* | — |
| SDNN | *Student* | *Pressure, Sex, Position* | — |
| HRVTi | *Student* | *Pressure, Sex, Position* | — |
| RMSDD | *Student* | *Pressure, Sex, Position* | — |
| BRS | *Student* | *Pressure, Sex, Position* | — |
| LFNorm | *Student* | *Pressure, Sex, Position* | — |
| HFNorm | *Student* | *Pressure, Sex, Position* | — |
| LF/HF | *Student* | *Pressure, Sex, Position* | *Pressure × Sex* |
| IOP | *Student* | *Pressure, Sex, Position, Side* | — |
| OPP | *Student* | *Pressure, Sex, Position, Side* | — |
| A<sub>IJV</sub> | *Student* | *Pressure, Sex, Position, Side* | — |
| IJVP | *Student* | *Pressure, Sex, Position, Side* | $\log(\sigma) \sim \text{Pressure}$ |
| IJVF<sup>§</sup> | *Cumulative* | *Pressure, Sex, Position, Side* | — |
| A<sub>CCA</sub> | *Student* | *Pressure, Sex, Position, Side* | — |
| PSV | *Student* | *Pressure, Sex, Position, Side* | — |
| EDV | *Student* | *Pressure, Sex, Position, Side* | *Pressure × Sex,*<br>*Sex × Position,*<br>*Sex × Side* |
| Age | — | — | — |
| Height | — | — | — |
| Weight | — | — | — |
| BMI | — | — | — |

Notes:
<sup>†</sup>Interactions or distributional parameters.
<sup>‡</sup>Bayes Factor analysis strongly favored a Gaussian distribution over a robust Student distribution for RPP.
<sup>§</sup>As described in the main text, IJVF flow pattern was modeled as an ordinal logistic regression with a logit link.
<sup>||</sup>Subject characteristics modeled as a unique group-level intercept.

Weakly informative priors were chosen across all parameters and they are summarized in Table 3. Normal priors were used for all *β* and *ζ*. Following the recommendations of Gelman^62^, half-Cauchy distributions were used for all *σ* and *σ_u_*. Gamma priors were used for all *ν*. The covariance matrix, **Σ**, was assigned a Lewandowski-Kurowicka-Joe (LKJ) prior^63,64^. Prior predictive checks were conducted to ensure that the priors generated credible estimates.

**Table 3:** Weakly informative priors used for the Bayesian multivariate dose-response model. Note that the priors apply to all relevant standardized dependent variables, *p*, in line with the model formulae outlined in Table 2.

| Group | Prior | Comment |
| --- | --- | --- |
| $\beta_0$ | $\mathcal{N}(0, 1)$ | Intercept |
| $\beta$ | $\mathcal{N}(0, 1)$ | Slope<sup>†</sup> |
| $\zeta_0$ | $\mathcal{N}(0, 1)$ | Distributional intercept parameter |
| $\zeta_{LBNP}$ | $\mathcal{N}(0, 1)$ | Distributional slope parameter |
| $\nu$ | $\Gamma(2, 0.1)$ | Degrees of freedom |
| $\sigma$ | $Cauchy^+(0, 1)$ | Population variance |
| $\sigma_u$ | $Cauchy^+(0, 1)$ | Group variance |
| $\mathbf{L}, \mathbf{R}$ | $\mathcal{LKJ}_{corr}(1)$ | Correlation structure<sup>‡</sup> |

Notes:
<sup>†</sup>Applies to $\beta_{LBNP}$ , $\beta_{sex}$ , $\beta_{pos}$ , $\beta_{side}$ (where relevant), and any interactions.
<sup>‡</sup>Lewandowski-Kurowicka-Joe (LKJ) Cholesky correlation distribution with shape parameter $\eta = 1$ .

The model was fit via Markov Chain Monte Carlo (MCMC) using Stan version 2.26.1^65^, R version 4.2.2^66^, and the brms package^67–69^. Stan is a probabilistic programming platform for statistical modeling and high-performance statistical computation, where Hamilton Monte Carlo (HMC) sampling is performed using a no-U-turn sampler (NUTS) to efficiently explore posteriors in models. The model was sampled using 20,000 draws (1,000 burn-in) in each of four chains. In the fitted model, chain diagnostics were visually inspected to ensure good mixing, with all 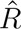 values *<* 1.01 and all effective chain lengths *>* 5000^70^. Posterior predictive checks were conducted to ensure that the posterior estimates approximated the data distribution. Pareto-smoothed leave-one-out cross validation was conducted to ensure accurate model predictive power^71^. All posterior summaries are given using the maximum a-posteriori estimate and the 89% highest density interval (HDI, also known as the 89% credible interval − 89% CrI). We further report evidence of existence and significance of effects, where existence is defined as *“the consistency of an effect in one particular direction (i.e., positive or negative), without any assumptions or conclusions as to its size, importance, relevance, or meaning”*^72^, and significance is *“being worthy of attention or important”*^72^. Evidence for the existence of an effect is presented using the probability of direction (*pd*), defined as the proportion of the posterior distribution that is of the same sign as its median (from 50% to 100%)^72^. Evidence for the significance of an effect (as an analogy to a frequentist *p*-value) is presented as the percentage of the full posterior for any particular effect (*Pressure*, *Sex*, *Position*, or *Side*) inside a region of practical equivalence (ROPE). For the majority of parameters, the ROPE is defined as [−0.05, 0.05] on a normalized scale (or [−0.1*sd*(*y_p_*), 0.1*sd*(*y_p_*)] on the original scale of measurement)^49,73^. For the log odds parameters related to ĲVF, the ROPE is defined as [−0.1*π/*√3, 0.1*π/*√3]^73^. In general, if more than 95% of the full posterior distribution is inside the ROPE, this can be interpreted as strong evidence in favor of the null hypothesis (no effect), whilst if less than 5% of the full posterior distribution is inside the ROPE, this can be interpreted as strong evidence of an effect^74^. Both *pd* and %*_ROPE_* must be interpreted together. For example, a posterior distribution that is centralized about 0 but very wide, may have a low %*_ROPE_*, whilst *pd* ≈ 50% (half the distribution is positive and half of the distribution is negative). This implies that, whilst there may be an effect, there is little evidence as to whether that effect is positive or negative.

## 3 Results

The raw data collected across all measured variables are presented and fully described in Appendix A. From the fitted dose-response model, we can extract and visualize the posterior draws for the effect size of each individual main effect (*Pressure*, *Sex*, *Position*, or *Side*), on each dependent variable *p*. Since all dependent variables in the model were standardized, the effect sizes can be compared across different variables. Sections 3.1, 3.2, 3.2, and 3.2 below consider the effect sizes of *Pressure*, *Sex*, *Position*, and *Side*, respectively, from the fitted dose-response curves.

Table 4 presents the fitted parameters from the dose-response curves, back-transformed from the standardized posterior distributions into their original units. Finally, Table 5 presents the *pd* and %*_ROPE_* for each of the four main effects (*Pressure*: *β_LBNP_*, *Sex*: *β_sex_*, *Position*: *β_pos_*, and *Side*: *β_side_*) for all dependent variables. *pd* and %*_ROPE_* are invariant of the scale used (normalized or original) since the ROPE scales with the dependent variable.

**Table 4:**
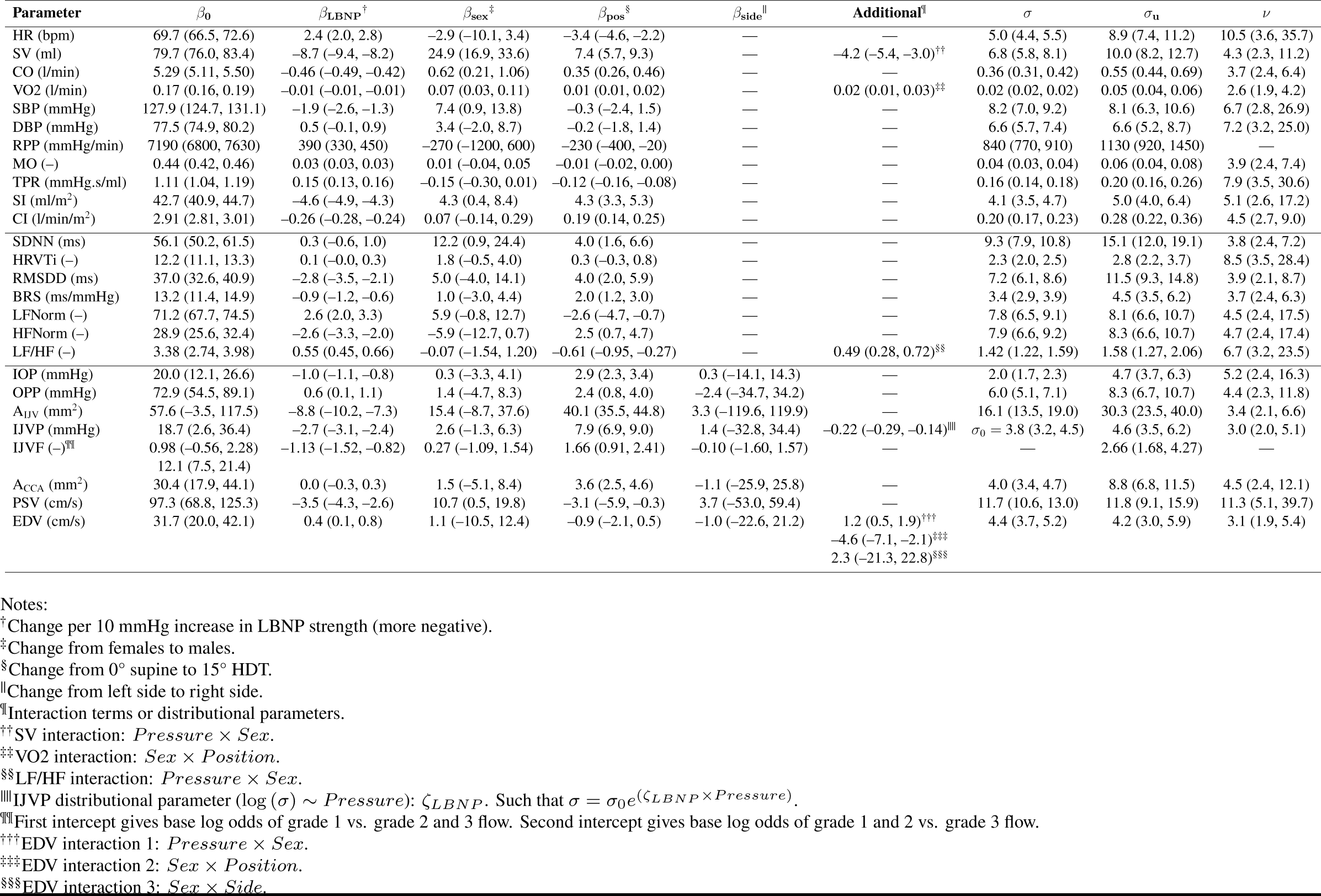
Posterior estimates for dose-response curves fitted to all measured parameters. Data are presented as the maximum a-posteriori (89% CrI).

**Table 5:** Summary of main effects in the multivariate dose-response model for all dependent variables considered. Evidence for existence of effects is presented as probability of direction (*pd*, where a higher % indicates more evidence for a consistent effect in a particular direction). Evidence for significance of effects is presented as percentage of full posterior distribution in region of practical equivalence (%*_ROPE_*, where less than 5% can be interpreted as strong evidence for an effect). See Section 2.5 for detail on the *pd* and ROPE range.

| Parameter | $\beta_{\text{LBNP}}$ | | $\beta_{\text{sex}}$ | | $\beta_{\text{pos}}$ | | $\beta_{\text{side}}$ | |
| --- | --- | --- | --- | --- | --- | --- | --- | --- |
| | $pd$ | $\%_{ROPE}$ | $pd$ | $\%_{ROPE}$ | $pd$ | $\%_{ROPE}$ | $pd$ | $\%_{ROPE}$ |
| HR | 100% | 0% | 77.83% | 19.41% | 100% | 0.30% | — | — |
| SV | 100% | 0% | 100% | 0% | 100% | 0% | — | — |
| CO | 100% | 0% | 98.95% | 2.76% | 100% | 0.01% | — | — |
| VO2 | 100% | 0.81% | 99.61% | 0.79% | 99.91% | 20.20% | — | — |
| SBP | 100% | 8.73% | 96.22% | 5.60% | 63.28% | 72.76% | — | — |
| DBP | 91.38% | 98.11% | 85.80% | 14.04% | 58.30% | 70.47% | — | — |
| RPP | 100% | 0% | 70.06% | 22.47% | 96.34% | 38.59% | — | — |
| MO | 100% | 0% | 65.67% | 23.91% | 97.20% | 30.62% | — | — |
| TPR | 100% | 0% | 93.69% | 10.02% | 100% | 0.05% | — | — |
| SI | 100% | 0% | 95.86% | 8.47% | 100% | 0% | — | — |
| CI | 100% | 0% | 70.72% | 30.82% | 100% | 0% | — | — |
| SDNN | 69.18% | 99.99% | 95.76% | 5.42% | 99.46% | 11.75% | — | — |
| HRVTi | 90.29% | 99.51% | 89.01% | 10.87% | 78.78% | 62.93% | — | — |
| RMSDD | 100% | 0.91% | 81.00% | 17.44% | 99.93% | 4.21% | — | — |
| BRS | 100% | 15.46% | 62.06% | 22.99% | 99.98% | 0.92% | — | — |
| LFNorm | 100% | 0.28% | 91.85% | 10.89% | 98.28% | 18.63% | — | — |
| HFNorm | 100% | 0.34% | 91.93% | 10.85% | 98.42% | 18.73% | — | — |
| LF/HF | 100% | 0% | 56.70% | 23.67% | 99.71% | 5.36% | — | — |
| IOP | 100% | 0% | 56.96% | 17.16% | 100% | 0% | 51.25% | 4.48% |
| OPP | 97.77% | 97.06% | 66.81% | 21.46% | 99.35% | 10.73% | 51.62% | 4.28% |
| A<sub>IJV</sub> | 100% | 0% | 85.52% | 13.18% | 100% | 0% | 51.39% | 4.47% |
| IJVP | 100% | 0% | 86.11% | 21.65% | 100% | 0% | 51.52% | 4.47% |
| IJVF | 100% | 0% | 61.21% | 16.74% | 100% | 0.04% | 50.10% | 14.53% |
| A<sub>CCA</sub> | 52.12% | 100% | 64.46% | 16.35% | 100% | 0% | 51.23% | 4.51% |
| PSV | 100% | 0.28% | 95.60% | 5.94% | 96.03% | 26.60% | 53.70% | 4.47% |
| EDV | 97.12% | 93.87% | 55.09% | 8.32% | 84.96% | 44.97% | 50.77% | 4.45% |

### 3.1 Pressure Effect

Figure 2 presents the normalized effect size responses of all variables considered with respect to the main effect *Pressure*. The variables are ordered from the largest positive effect size at the top of the figure to the largest negative effect size at the bottom of the figure. In addition, ĲVF is presented separately due to the differing ROPE range.

**Figure 2:**
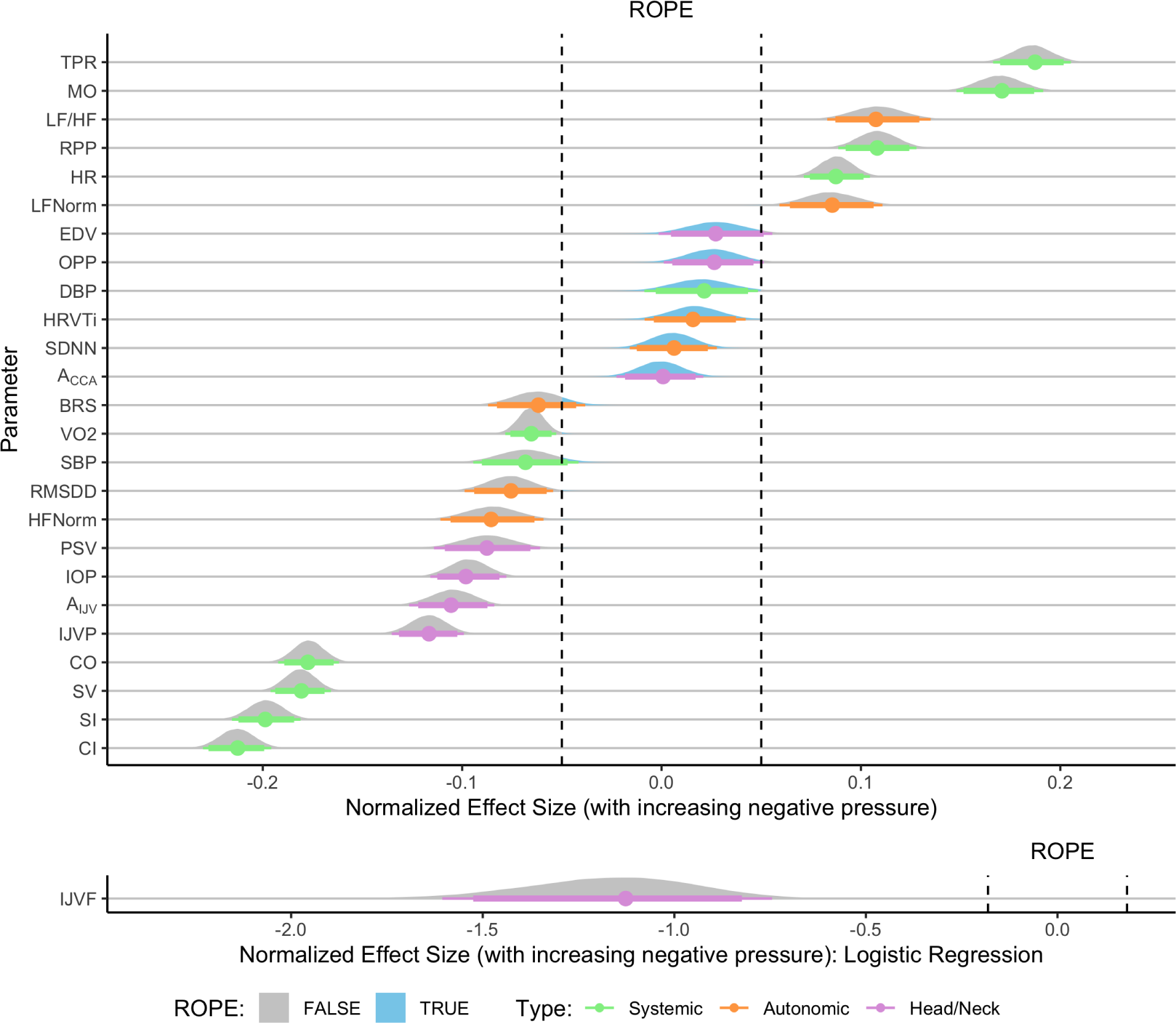
Normalized effect size responses of systemic (green), autonomic (orange), and head/neck (purple) variables to the main effect LBNP *Pressure*. Variables are ordered from the largest positive effect size at the top of the figure to the largest negative effect size at the bottom of the figure. Data are presented as the posterior distributions from the Bayesian multivariate regression model. Distributions are colored gray when located outside of the ROPE, and they are colored blue when located inside the ROPE. Points and error bars underneath the distributions represent the maximum a-posteriori estimate along with the 89% (thick) and 95% (thin) highest density intervals. ĲVF is presented separately below, since the ROPE is defined differently for a logistic regression model (see Section 2.5 for detail).

Of the four main effects considered, the effect of LBNP *Pressure* presents the strongest evidence (i.e., narrowest posterior distributions), with the majority of parameters’ posterior distributions situated either fully inside or outside of the ROPE. The systemic hemodynamic variables are generally those that are most influenced by LBNP *Pressure*, with a larger relative effect size (either positive or negative) than the autonomic or cephalad variables. In particular, we observe a decrease in SV/CO (and their indexed equivalents) with increasing negative pressure, as well as the corresponding rise in TPR and MO. In contrast, there is strong evidence in favor of no effect of LBNP *Pressure* on five variables (A_CCA_, SDNN, HRVTi, DBP, and OPP) and trending evidence of no effect in EDV (%*_ROPE_*: 93.87%). These results suggest that a) overall heart rate variability is not influenced by LBNP, and b) most LBNP-related effects are related to the systolic (%*_ROPE_* = 8.73%), as opposed to the diastolic (%*_ROPE_* = 98.11%), part of the blood pressure waveform. In addition, related to the head/neck hemodynamics, it is insightful that there is strong evidence for a significant effect of LBNP on IOP, A_ĲV_, and ĲVP (%*_ROPE_*: 0% for all three) of approximately similar relative magnitude, yet no effect on OPP (%*_ROPE_*: 97.06%). This has potential implications for the use of LBNP as a SANS countermeasure, discussed in Section 4 below.

Of the three groups (systemic hemodynamics, autonomic response, and head/neck), the autonomic variables are the least affected by LBNP, although there is still strong evidence of a decrease in parasympathetic activity (decrease in RMSDD and HFNorm; %*_ROPE_*: 0.91% and 0.34%, respectively) matched by an increase in sympathetic activity (increase in LFNorm and LF/HF; %*_ROPE_*: 0.28% and 0%, respectively). Finally, there is a clear effect of LBNP on ĲVF flow pattern (%*_ROPE_*: 0%), with the relative log odds of a higher grade (2 or 3) flow decreasing by 1.13 (89% CrI: 0.82 to 1.52) with each 10 mmHg increase in LBNP strength.

### 3.2 Sex Effect

Figure 3 presents the normalized effect size responses of all variables considered with respect to the main effect *Sex*. The variables are ordered from the largest positive effect size at the top of the figure to the largest negative effect size at the bottom of the figure. In addition, ĲVF is presented separately due to the differing ROPE range. Due to the contrast coding in the model, a positive effect size in a variable represents an increase in that variable in male subjects compared to female subjects. Tables 4 and 5 present the fitted parameters from the dose-response curves, and the main effects *pd* and %*_ROPE_*, respectively.

**Figure 3:**
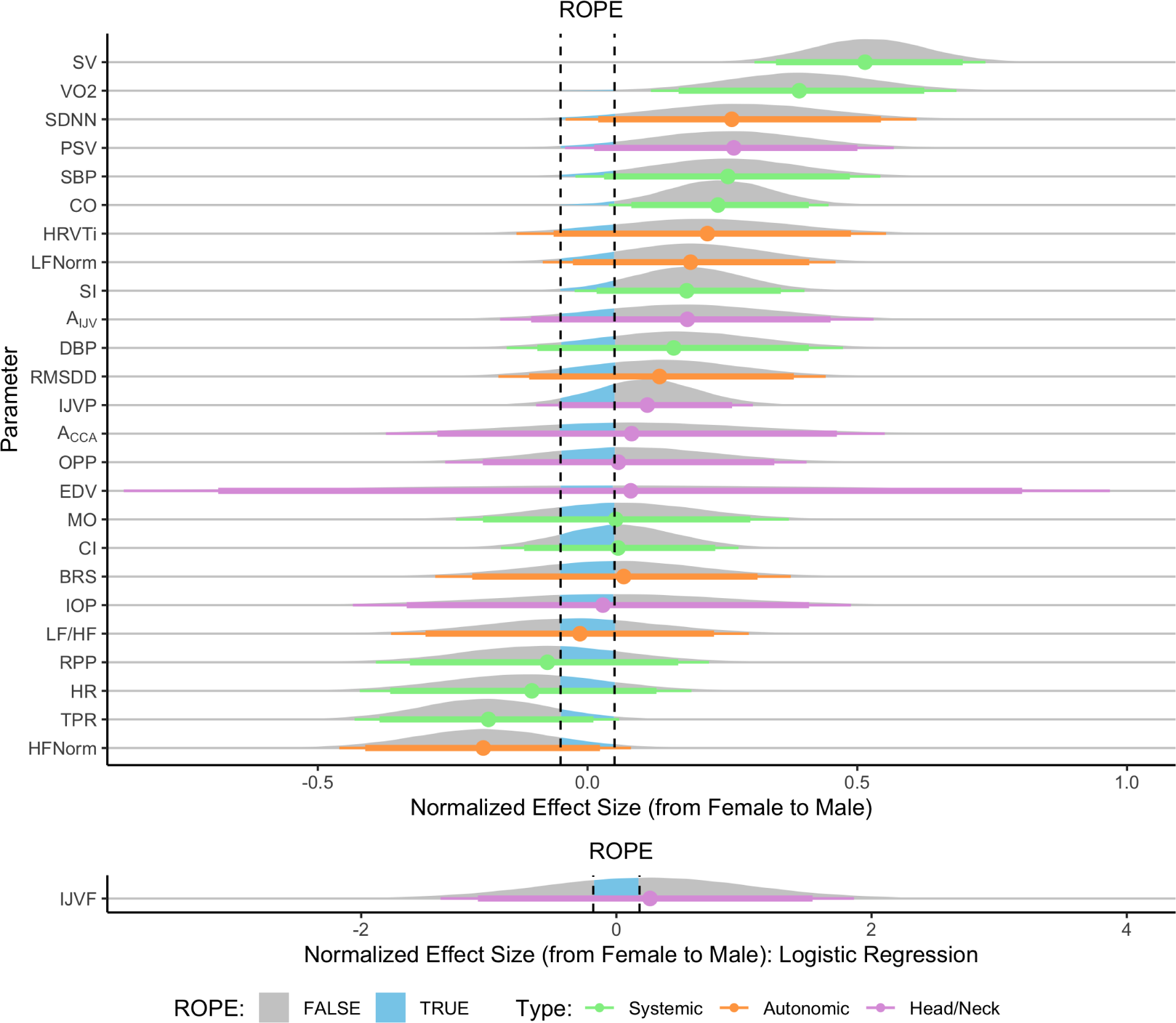
Normalized effect size responses of systemic (green), autonomic (orange), and head/neck (purple) variables to the main effect *Sex* (male or female). A positive effect size represents an increase in male subjects with respect to female subjects. Variables are ordered from the largest positive effect size at the top of the figure to the largest negative effect size at the bottom of the figure. Data are presented as the posterior distributions from the Bayesian multivariate regression model. Distributions are colored gray when located outside of the ROPE, and blue when located inside the ROPE. Points and error bars underneath the distributions represent the maximum a-posteriori estimate along with the 89% (thick) and 95% (thin) highest density intervals. ĲVF is presented separately below, since the ROPE is defined differently for a logistic regression model (see Section 2.5 for detail).

In contrast to the effect of *Pressure*, the posteriors associated with the *Sex* effect are much wider. This is due to the fact that the magnitude of any sex effect between males and females is often overshadowed by the natural inter-subject variability found across all subjects. We found strong evidence of significant effects of *Sex* in only three variables (SV, %*_ROPE_* = 0%; VO2, %*_ROPE_* = 0.79%; and CO, %*_ROPE_* = 2.76%). With respect to the indexed variables, there is minimal evidence of a sex effect in CI (*pd* = 70.72%, %*_ROPE_* = 30.82%) but some evidence of a sex effect in SI (*pd* = 95.86%, %*_ROPE_* = 8.47%). The presence of a significant effect in the absolute variables (CO, SV), and the lack of a significant effect in the indexed variables (and elsewhere) would appear to indicate that differences between males and females are principally driven by anthropometric differences (i.e., on average males are larger, with a higher total blood volume and a larger stroke volume).

In five variables, there is moderate evidence of the presence of a sex effect, even if it is not necessarily of a significant magnitude. This is evidenced by variables with a *pd* greater than 90% and a %*_ROPE_* less than 10% (in simple terms, the majority of the posterior distribution is away from 0). These variables are SDNN (*pd* = 95.76%, %*_ROPE_* = 5.42%), PSV (*pd* = 95.60%, %*_ROPE_* = 5.94%), SBP (*pd* = 96.22%, %*_ROPE_* = 5.60%), SI (*pd* = 95.86%, %*_ROPE_* = 8.47%), and TPR (*pd* = 93.69%, %*_ROPE_* = 10.02%). The fact that SDNN appears larger in males (by, on average, 12.2 ms, 89% CrI: 0.9 to 24.4 ms) is likely explained by hormonal effects on the autonomic nervous system^75^. We do not, however, find a corresponding reduction in RMSDD in males, which would be indicative of higher parasympathetic activity in females.

In summary, comparison between absolute variables (CO, SV) and indexed variables (CI, SI) suggest that sex differences in the data appear to be principally driven by anthropometric variation between males and females. However, there is some evidence of increased sympathetic activity in male subjects.

### 3.3 Position Effect

The main effect of position is insightful as it relates to the characterization of LBNP responses in the presence of a cephalad fluid shift. Figure 4 presents the normalized effect size responses of all variables considered with respect to the main effect *Position*. The variables are ordered from the largest positive effect size at the top of the figure to the largest negative effect size at the bottom of the figure. In addition, ĲVF is presented separately due to the differing ROPE range. Based on the coding scheme that was used to capture the main effect *Position* in the model, a positive effect size represents an increase from 0° supine to 15° HDT. Tables 4 and 5 present the fitted parameters from the dose-response curves, and the main effects *pd* and %*_ROPE_*, respectively.

**Figure 4:**
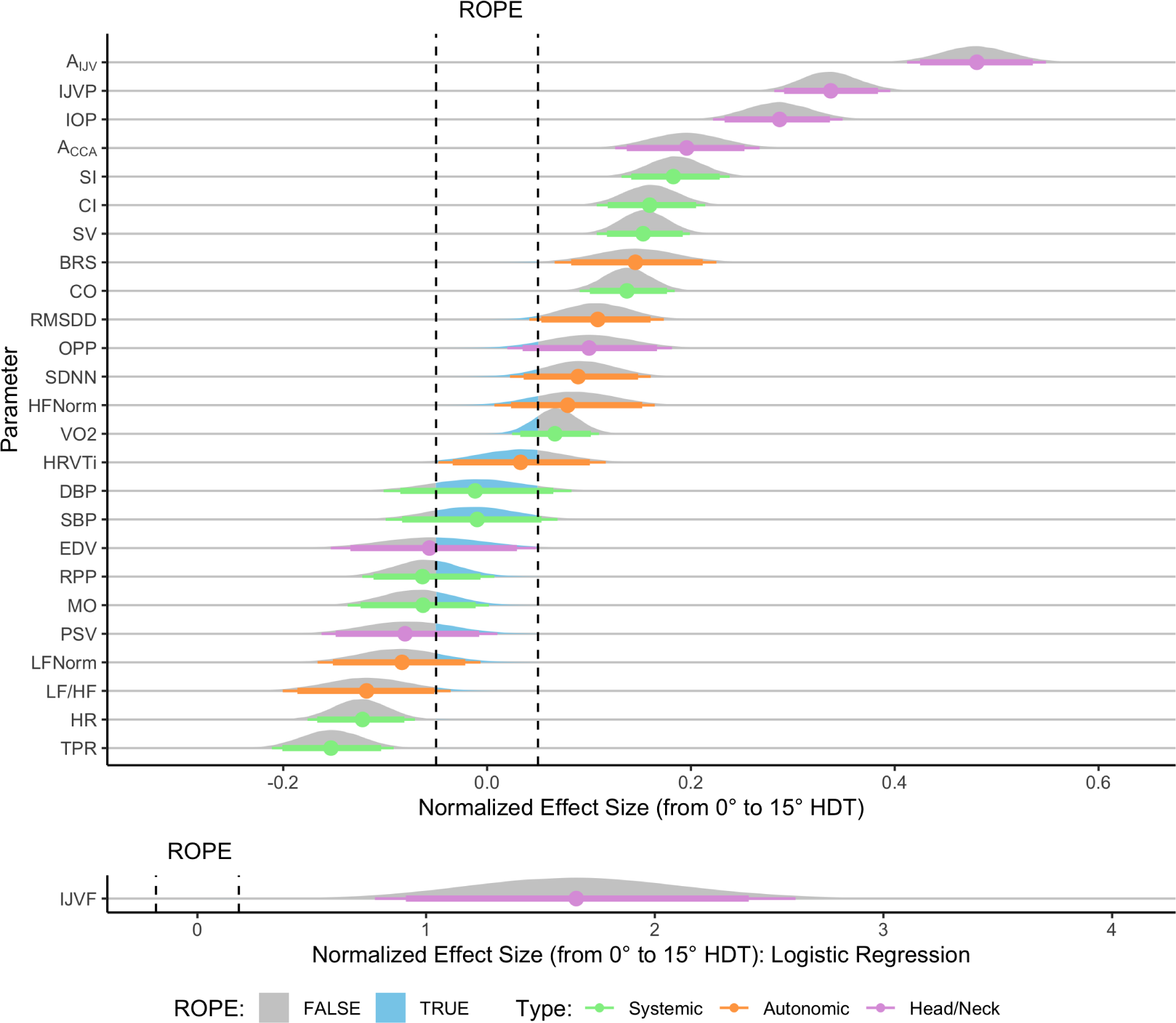
Normalized effect size responses of systemic (green), autonomic (orange), and head/neck (purple) variables to the main effect *Position* (0° supine or 15° HDT). A positive effect size represents an increase in 15° HDT with respect to 0° supine. Variables are ordered from the largest positive effect size at the top of the figure to the largest negative effect size at the bottom of the figure. Data are presented as the posterior distributions from the Bayesian multivariate regression model. Distributions are colored gray when outside of the ROPE, and blue inside the ROPE. Points and error bars underneath the distributions represent the maximum a-posteriori estimate along with the 89% (thick) and 95% (thin) highest density intervals. ĲVF is presented separately below, since the ROPE is defined differently for a logistic regression model (see Section 2.5 for detail).

A significant effect is seen in six of the 11 systemic hemodynamic parameters, two of the autonomic parameters, and five of the eight head/neck parameters. The largest effect sizes are found in the jugular vein response (area, pressure, and flow). This is congruent with our previous studies^76^, which demonstrated the strong gravitational dependence of the jugular vein. In contrast to Whittle and Diaz-Artiles^76^, we also find a significant effect of Position on A_CCA_ (*pd* = 100%, %*_ROPE_* = 0%), although it should be noted that in the present study we are only considering a unique tilt angle. We also note significant increases with 15° HDT in SV (*pd* = 100%, %*_ROPE_* = 0%), CO (*pd* = 100%, %*_ROPE_* = 0%) (and their indexed equivalents), IOP (*pd* = 100%, %*_ROPE_* = 0%), BRS (*pd* = 99.98%, %*_ROPE_* = 0.92%), and RMSDD (*pd* = 99.93%, %*_ROPE_* = 4.21%), along with significant decreases in HR (*pd* = 100%, %*_ROPE_* = 0.30%) and TPR (*pd* = 100%, %*_ROPE_* = 0.05%).

The RMSDD and BRS responses, combined with the decrease in HR, indicate that the autonomic response is activated by the cephalad fluid shift, manifested principally by an increase in vagal activity lowering HR. This is combined with the reduced TPR promoting venous return, leading to increased SV and CO through the Frank-Starling mechanism^24^.

Finally, we note an increase in OPP in 15° HDT (*pd* = 99.35%, %*_ROPE_* = 10.73%), which is significant in direction and approaching significance in magnitude. This is congruent with our previous studies demonstrating an increase in OPP in HDT^57^. Analysis of the relative magnitudes of the effect sizes indicates that the increase in OPP is blunted by the constancy of blood pressure, both SBP and DBP, which present evidence in favor of the null hypothesis of no effect of position (SBP: *pd* = 63.28%, %*_ROPE_* = 72.76%; DBP: *pd* = 58.30%, %*_ROPE_* = 70.47%).

### 3.4 Side Effect

Figure 5 presents the normalized effect size responses of the head/neck variables with respect to the main effect *Side*. The variables are ordered from the largest positive effect size at the top of the figure to the largest negative effect size at the bottom of the figure. In addition, ĲVF is presented separately due to the differing ROPE range. Due to the contrast coding in the model, a positive effect size represents an increase in the right side with respect to the left side. Tables 4 and 5 present the fitted parameters from the dose-response curves, and the main effects *pd* and %*_ROPE_*, respectively.

**Figure 5:**
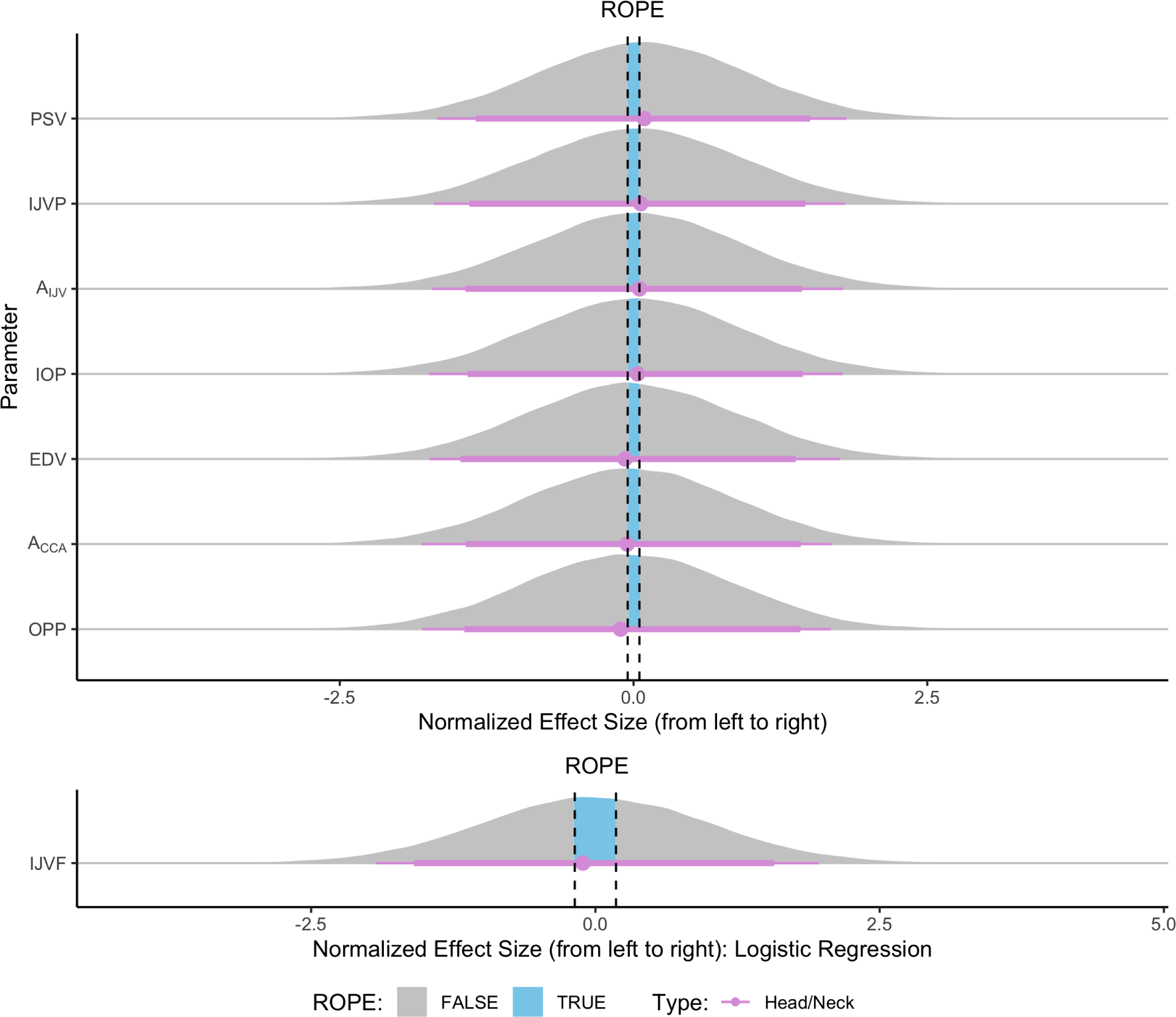
Normalized effect size responses of head/neck (purple) variables to the main effect *Side* (left or right). A positive effect size represents an increase in the right side with respect to the left side. Variables are ordered from the largest positive effect size at the top of the figure to the largest negative effect size at the bottom of the figure. Data are presented as the posterior distributions from the Bayesian multivariate regression model. Distributions are colored gray when outside of the ROPE, and blue inside the ROPE. Points and error bars underneath the distributions represent the maximum a-posteriori estimate along with the 89% (thick) and 95% (thin) highest density intervals. ĲVF is presented separately below, since the ROPE is defined differently for a logistic regression model (see Section 2.5 for detail).

In all eight variables considered, we did not find an effect of *Side*. For all variables except ĲVF, both the %*_ROPE_* is small (*<* 5%), and the *pd* is around 50% to 54%. This is due to the broad posterior distributions roughly centered around 0. Thus, there is no evidence of an effect size in any particular direction. This is in contrast to our results from previous tilt experiments^57,76,77^, where we found differences in size between the left and right internal jugular veins. It would appear that this side effect is only present in the amplified expansion of the internal jugular vein in large head-down tilts. Note that, in the Bayesian framework, our results suggest that there is no significant evidence either in favor of the null hypothesis (no effect), or the alternative (an existing effect). Aside from our findings on A_ĲV_ in tilt discussed above, the rest of the results presented here are congruent with our previous results, such that we found no effect of side on IOP, OPP, or ĲVP.

### 3.5 Internal Jugular Vein Flow

The majority of the variables are explained by linear models, whose interpretation is relatively straight forward. In contrast, the dose-response curve for the internal jugular vein blood flow velocity waveform pattern is based on an ordinal logistic regression, with a slightly more cryptic interpretation. Thus, it is insightful to discuss this dose-response in detail. Figure 6 presents the ĲV flow pattern dose-response curve. Based on the effect sizes in Figures 2, 3, 4, and 5 (where there was evidence of *Pressure* and *Position* effects, but no evidence of *Sex* or *Side* effects), we have grouped males and females, and left and right sides together. Whilst the response is ordinal, the figure shows the latent variable given by the logit function^78^. Due to only four instances of Grade 3 flow occurring (Figure A4), we have grouped the probabilities such that the Y axis in the curve represents the probability of “greater than Grade 1 flow” (i.e., Grade 2 or Grade 3 flow).

**Figure 6:**
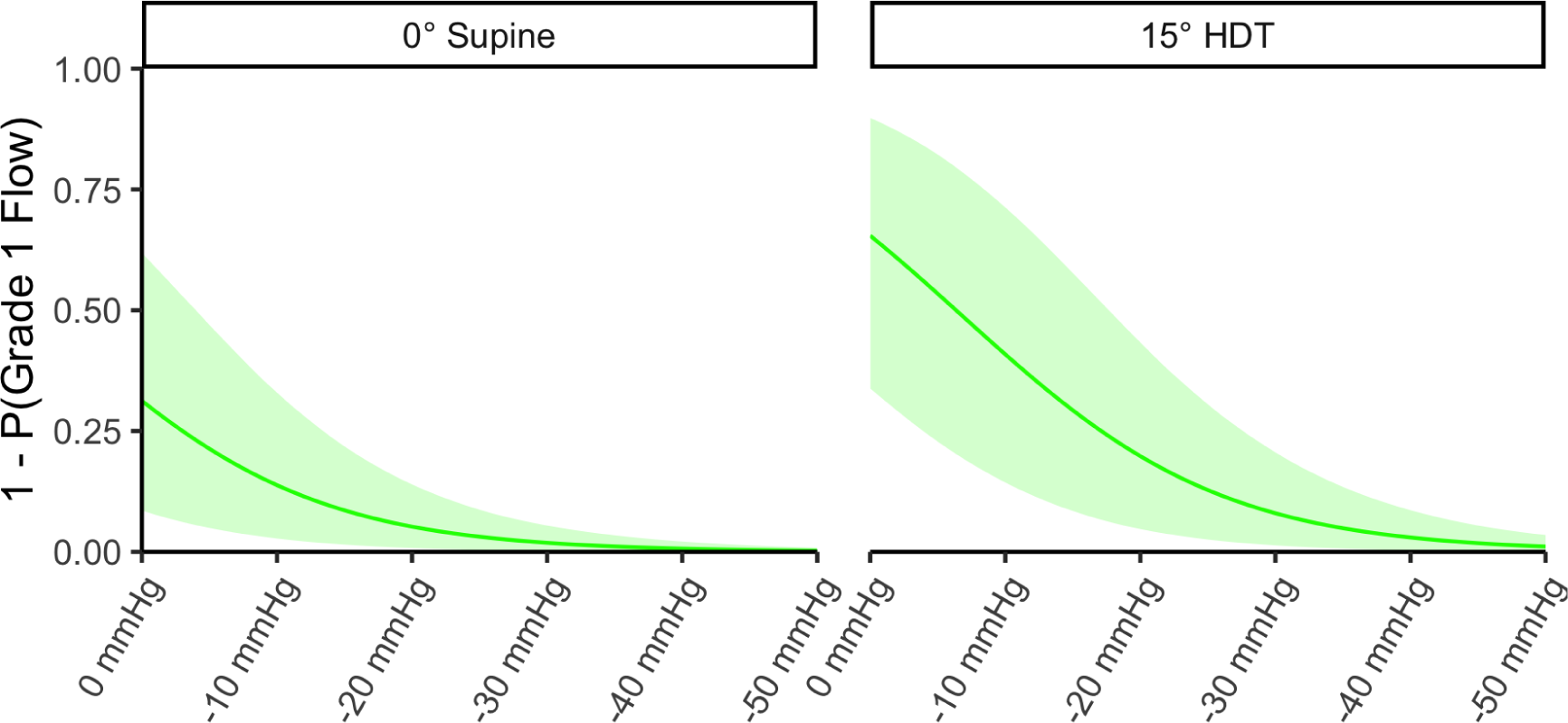
Dose-response curve for internal jugular vein blood flow velocity waveform pattern. The *y*-axis represents the probability of having a higher than grade 1 flow (i.e., grade 2 or grade 3 flow). Dose-response represents the fitted posterior draws from the multivariate regression model. *Sex* and *Side* effects are pooled. *Position* (0° supine or 15° HDT) is faceted. Dose-response is presented as the maximum a-posteriori estimate ± 89% CrI.

In 0° supine, the probability of greater than Grade 1 flow is 31.2% (89% CrI: 8.5% to 61.9%) at 0 mmHg. This is reduced by LBNP to 13.8% (89% CrI: 2.8% to 33.0%), 5.2% (89% CrI: 0.8% to 13.9%), 1.9% (89% CrI: 0.2% to 5.5%), 0.7% (89% CrI: 0.1% to 2.1%), and 0.2% (89% CrI: 0.0% to 0.8%) at −10 mmHg, −20 mmHg, −30 mmHg, −40 mmHg, and −50 mmHg, respectively. In 15° HDT, there is an increased probability of Grade 2 or higher flow of 65.4% (89% CrI: 33.8% to 89.8%) at 0 mmHg. This is reduced by LBNP to 40.9% (89% CrI: 14.3% to 71.3%), 19.8% (89% CrI: 4.7% to 43.3%), 8.0% (89% CrI: 1.3% to 20.6%), 3.0% (89% CrI: 0.3% to 8.6%), and 1.1% (89% CrI: 0.1% to 3.5%) at −10 mmHg, −20 mmHg, −30 mmHg, −40 mmHg, and −50 mmHg, respectively.

The data highlight that LBNP effectively increases blood flow in the internal jugular veins, particularly in 15° HDT. The implications of these results will be further discussed in Section 4 below.

### 3.6 Multivariate Relationships

Figure 7 presents the multivariate relationships amongst all of the variables considered, including the subject characteristics (*Age*, *Height*, *W eight*, and *BMI*). Correlations are only displayed where there is significant evidence of an effect (in the Bayesian formulation, this is when the full posterior distribution does not encompass 0)ii. The inclusion of the subject characteristics also provides insight into any relationships between the variables driven by anthropometric considerations such as height or weight.

**Figure 7:**
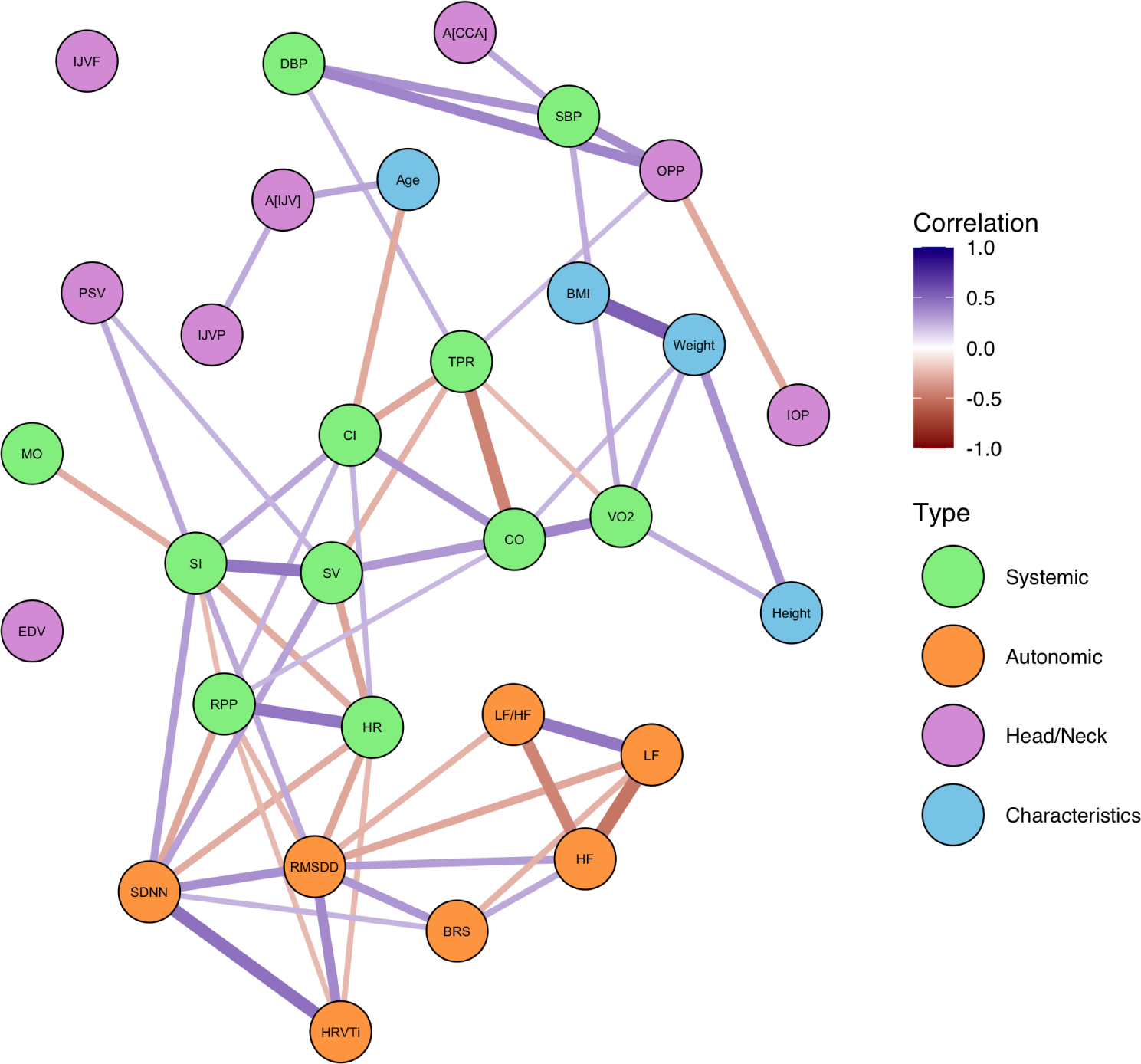
Graph structure representing the multivariate relationships amongst all of the measured variables (green: systemic, orange: autonomic; purple: head/neck; blue: subject characteristics). The direction of the correlations (positive or negative) are represented by the color of the edges, and the strength of the correlations (the maximum a-posteriori estimate) are represented by the thickness of the edges. Only significant correlations are shown (see text for details).

Broadly, the graph structure appears to form two connected groups. All of the autonomic parameters (in orange) are strongly connected to one another, forming one group. Similarly, the systemic hemodynamic parameters (in green) form a second connected group structure. HR is linked to the autonomic parameters through heart rate variability (associated with both SDNN and HRVTi) and parasympathetic activity (RMSDD), where higher HR is associated with lower variability and lower vagal activity. Similarly, increased HRV is associated with increased SV, and reduced RPP.

OPP is associated with both blood pressure numerics (SBP and DBP) and IOP, but increased OPP is also associated with increased TPR. The relationship with the subject characteristics is also insightful: in contrast to Buckey *et al.*^26,79^, we do not find any association between IOP and body weight or BMI. However greater height and weight are associated with increased oxygen consumption (VO2). More interestingly, we see that Age is positively correlated with ĲV cross-sectional area, A_ĲV_, (and by extension pressure, ĲVP), and negatively correlated with CI. Finally, carotid hemodynamics are associated with SBP (correlated with A_CCA_) and SV (correlated with PSV). There is no significant relationship between EDV or ĲVF and any of the other metrics.

## 4 Discussion

This study investigated the acute effects of LBNP on the cardiovascular system. To our knowledge, this is the most comprehensive analysis of cardiovascular hemodynamics, autonomic, and cephalad response to LBNP to date. Our main findings suggest that: (1) there is a varying magnitude of the normalized effect sizes of responses to LBNP in different cardiovascular variables; (2) sex differences exist between the male and female response; however, these appear to be principally driven by anthropometric considerations; (3) concerning head/neck variables (i.e., common carotid arteries, internal jugular veins, and eyes), there is no evidence of a difference between the response in the left and right sides to LBNP (within the experimental conditions considered); and (4) there is an underlying multivariate structure with associations connecting all but two (EDV, ĲVF) of the variables considered, including cardiovascular variables as well as subject characteristics such as age, height, weight, and BMI.

Multiple studies have investigated the effects of LBNP on cardiovascular hemodynamics; many of these are summarized in a comprehensive review article by Goswami *et al.*^17^. In a classic study by Blomqvist and Stone^80^, the systemic hemodynamic responses to graded LBNP up to −40 mmHg were shown to be linear for TPR, HR, SV, CO, and MAP. Data from their study were compiled by Goswami *et al.*^17^. Our study, and in particular Figure 2, extends this work and increases the number of variables considered. Our findings agree with the authors as to the relative magnitudes of the effects of TPR (large positive), HR (small positive), and CO and SV (large negative). We further find no decrease in DBP, and a small decrease in SBP, which correspond well with the minimal decrease in MAP found by Goswami and colleagues. In Figure 2 we add the effect sizes of 17 new variables, and present the relative magnitudes of the LBNP-induced changes. Of note, we find that, whilst the TPR and SV/CO changes are the largest positive and negative effects, respectively, there are smaller, still significant, negative effects surrounding the jugular vein (A_ĲV_, ĲVP, ĲVF), IOP, and VO2, as well as positive effects on myocardial oxygen supply:demand index (MO) and rate pressure product (RPP).

Murray *et al.*^81^ investigated graded LBNP in four supine subjects and found a linear increase in HR from 59 ± 2.9 bpm at 0 mmHg to 90 ± 5.5 bpm at −50 mmHg. They further found changes in blood pressure (SBP / DBP) from 125 ±6.7 / 71 ± 4.6 mmHg at 0 mmHg to 119 ± 4.3 / 81 ± 3.5 mmHg at −50 mmHg, and a decrease in SV and CO from 84 ± 6.0 ml and 4.9 ± 0.3 l/min at 0 mmHg to 39 ± 4.6 ml and 3.4 ± 0.2 l/min at −50 mmHg, respectively^81^. Murray’s increase in HR (31 bpm) is greater than the average increase in our study (12 bpm, 89% CrI: 10 to 14 bpm) over the same range, considering both positions and both sexes. Our SBP decreased at a similar rate, from 129.5 ±2.2 mmHg at 0 mmHg to 117.2 ± 2.4 mmHg at −50 mmHg, but we did not find a significant increase in DBP (effect size of 0.5 mmHg, 89% CrI: −0.1 to 0.9 mmHg per 10 mmHg LBNP). Conversely, we found a larger decrease in SV and CO (effect sizes: −0.87 ml/mmHg, 89% CrI: −0.94 to −0.82 ml/mmHg; and −0.046 l/min/mmHg, 89% CrI: −0.049 to −0.042 l/min/mmHg, respectively). In general, our values fall within the confidence intervals found by Murray *et al.*. Further, this study by Murray is one of the few to investigate systemic vascular resistance or TPR. They found an increase from 1,415 ± 123 dyne-sec.cm^−5^ at 0 mmHg to 2,200 ± 132 dyne-sec.cm^−5^ at −50 mmHg (equivalent to 1.06 ± 0.09 mmHg.s/ml to 1.65 ± 0.10 mmHg.s/ml in peripheral resistance units, PRU). This closely matches the increase we found, from 1.11 ± 0.04 mmHg.s/ml at 0 mmHg to 1.75 ± 0.06 mmHg.s/ml at −50 mmHg; an effect size of 0.015 mmHg.s/ml/mmHg (89% CrI: 0.013 to 0.016 mmHg.s/ml/mmHg). Similarly, Levine *et al.* found an increase of 18 bpm in HR, and a decrease of 43 ml in SV at −40 mmHg LBNP (with respect to 0 mmHg) in 13 supine subjects^82^. Finally, Hinojosa-Laborde *et al.* considered graded LBNP as an experimental model of hemorrhage^18^. They found a decrease in SBP from 121 to 80 mmHg, a decrease in SV from 59 to 41 ml, a decrease in CO from 5.1 to 4.2 l/min, and an increase in HR from 82 to 94 bpm from 0 mmHg to −40 mmHg LBNP. The decrease in SBP is much larger than the corresponding reduction in our data; however, all other measurements are within the calculated confidence intervals of the dose-responses.

Autonomic response to LBNP is principally mediated through the reduction in central blood volume lowering systemic flow and perfusion pressure, leading to stimulation of the arterial baroreflex^17,83–85^. Convertino *et al.* investigated the effect of LBNP on baroreflex sensitivity, finding that BRS decreased from 15 ± 1 ms/mmHg to 7 ± 1 ms/mmHg at presyncope in low-tolerance subjects and from 17 ± 2 ms/mmHg to 4 ± 0 ms/mmHg in high-tolerance subjects^86^. This matches well with our data, which show a reduction of BRS from 13.8 ± 1.1 ms/mmHg at 0 mmHg to 9.2 ± 0.7 ms/mmHg at −50 mmHg (average of both sexes and both positions). We did not deliberately take our subjects to the point of presyncope, but the decreasing trend is anticipated to continue to that point. This decrease in BRS, combined with the reduction in time- and frequency-derived HRV metrics (i.e., RMSDD and HFNorm) is indicative of progressive vagal withdrawal^87–89^. This vagal withdrawal is matched by a linear increase in sympathetic nervous activation with progressive LBNP. Thus, our data demonstrate a linear increase in LFNorm, matched by a linear increase in LF/HF. These data are supported by multiple studies assessing either HRV metrics or muscle sympathetic nervous activity (MSNA) with LBNP^28,35,36,90–92^. Experiments using cholinergic blockade demonstrate the importance of both arms of the autonomic response (vagal withdrawal and sympathetic activation) to mediate cardiac function in LBNP^93,94^. In addition, the sympathetic response is also important for mediating vascular smooth muscle constriction in response to the reduction in central blood volume^86^.

Regarding sex differences, Convertino investigated differences in autonomic function related to blood pressure regulation^37^. Our results are congruent with Convertino, who noted higher HR in female subjects, combined with a lower SV during LBNP. Convertino took all subjects to presyncope, and noted a lower tolerance in females. Whilst we did not deliberately take our subjects to presyncope, the fact that six of the seven subjects who reached presyncope whilst supine in the 0 to −50 mmHg range were female suggests a lower orthostatic tolerance in females. This finding is well supported by multiple studies^29,37,39,40,95,96^. Convertino further derived the effect size of the LBNP response in males and females with regard to HR, SV, CO, MAP, and TPR, noting that the difference in slope between males and females was non-significant (*p >* 0.05) in all cases except for TPR (*p* = 0.0002). For HR, he noted a 0.37 ± 0.05 bpm/mmHg increase in males and a 0.58 ± 0.10 bpm/mmHg increase in females. This is slightly higher than the 0.24 bpm/mmHg (89% CrI: 0.20 to 0.28 bpm/mmHg) that we derived. With regards to SV, Convertino found a −1.06 ± 0.10 ml/mmHg change in males and a −1.23 ± 0.19 ml/mmHg change in females. This compares with our findings of −0.87 ml/mmHg (89% CrI: −0.94 to −0.82 ml/mmHg). In contrast, we also note evidence for the existance of an interaction effect, with males decreasing SV 0.42 ml/mmHg (89% CrI: 0.30 to 0.54 ml/mmHg) faster than females. Finally, in CO, we note a change of −0.046 l/min/mmHg (89% CrI: −0.049 to −0.042 l/min/mmHg). This fits in between Convertino’s measured values of −0.03 ± 0.01 l/min/mmHg in males and −0.07 ± 0.01 l/min/mmHg in females. On the autonomic side, Convertino measured the baroreflex sensitivity (BRS), noting that the response was 1.32 ms/mmHg lower in females (*p* = 0.047). We found that BRS was potentially higher in males by 1.0 ms/mmHg (89% CrI: −3.0 to 4.4 ms/mmHg), though the difference was far from significant and outweighed by intersubject variability (*pd* = 62.06%, %*_ROPE_* = 22.99%). Other studies have found potential differences in the autonomic response between men and women. Frey *et al.* and Evans *et al.* found that women have a more dominant vagal response, whilst men primarily demonstrate a greater sympathetic response^34,97^. In addition, Frey and Hoffler also found that men exhibited a larger increase in TPR^38^. By considering Figure 3, we likewise find evidence of a higher sympathetic response in males based on LFNorm (*pd* = 91.85%, %*_ROPE_* = 10.89%); however, we find conflicting evidence related to a higher vagal response in women. The two autonomic markers corresponding to vagal response trend in opposite directions, with HFNorm higher in females (*pd* = 91.93%, %*_ROPE_* = 10.85%), but RMSDD higher in males (*pd* = 81.00%, %*_ROPE_* = 17.44%). Further study is required to investigate these discrepancies in the sex-driven autonomic response. More recently, Patterson *et al.* measured A_ĲV_ differences between males and females during LBNP exposure in the range 0 to −40 mmHg^41^. They noted a significant effect of sex (*p <* 0.001) and LBNP (*p <* 0.001), but no significant interaction (*p* = 0.066), with A_ĲV_ being larger in female subjects at 0 mmHg, −20 mmHg, and −30 mmHg. In contrast, we found a significant effect of LBNP of −0.88 mm^2^/mmHg (89% CrI: −1.02 to −0.73 mm^2^/mmHg) but no significant effect of sex. In fact, in our study we found trending evidence of a higher A_ĲV_ in males by 15.4 mm^2^ (89% CrI: −8.7 to 37.6 mm^2^; *pd* = 85.52%, %*_ROPE_* = 13.18%). Our results match previous studies, for example Jeon *et al.* and Magnano *et al.*, who found no difference in A_ĲV_ between males and females^98,99^.

The article by Magnano *et al.* is particularly interesting in that they note a positive association between A_ĲV_ and age in over 1000 subjects^99^. Using a totally different methodology, we also discovered this association, captured in Figure 7. This finding lends support both to their conclusions and also to our Bayesian modeling workflow. Magnano and colleagues hypothesize that the increased A_ĲV_ is linked to inhibited central venous drainage as a result of raised intra-abdominal pressure with increased BMI, which trends higher in older individuals. By contrast, although admittedly in a far smaller study, we find no direct link between BMI (or weight) and A_ĲV_. This suggests that the association might be related to other factors outside of BMI. Magnano *et al.* posit on the role of endothelial progenitor cells (EPCs) in the vascular remodeling process, noting sex differences related to pregnancy hormones^100^. Given evidence that EPCs decrease with increasing age^101^, it could be possible that EPCs play a role in the age-related differences in A_ĲV_. Continuing our discussion of Figure 7, we also note the negative association between age and CI, which is supported by a number of studies^102,103^. Further, we also find an association between OPP and TPR. It is difficult to find evidence in the literature for similar relationships, although Fındıkoğlu *et al.* noted a decrease in OPP in 34 subjects after hot-water immersion, which was also matched by a decrease in TPR^104^. Samsudin *et al.* noted a relationship between OPP and EDV (which we did not find), but noted no relationship between OPP and resistive index^105^. In this study, resistive index (RI) referred only to the specifics of the ocular vasculature and not on the TPR; thus, it is difficult to draw comparisons. The effect of position is important since it allows us to determine the relative magnitude of the effect of a headward fluid shift induced by HDT (and, by extrapolation, microgravity) compared to the LBNP effect. Of the three groups of variables (systemic, autonomic, and cephalad), we note that the largest effect size increases occur in the head/neck parameters including A_ĲV_, ĲVP, IOP, and ĲVF. Interestingly, in this experiment we also found evidence of an increase in A_CCA_, which we did not observe in our previous tilt study^76^. It should be noted that the present study examines only a single tilt angle, and compensatory mechanisms may prevent this increase from progressing at more severe HDT angles. Aside from A_CCA_, there are no data that contradicts our previous experiments, including evidence that blood pressure is maintained in tilt^57,76,77^.

We did not find any differences in any of the head/neck variables between the left and right side. This is supported by our previous work examining eye pressures in a tilt paradigm, where results suggest that there is no difference in the pressures in the ocular system between the left and right side (neither in IOP, nor in OPP)^57^. Further, in Whittle and Diaz Artiles (2023)^76^, we considered the gravitational effects of the carotid arteries and jugular veins in tilt. In that study, we did not measure PSV or EDV; however, we could anticipate that there is little difference between sides in those variables since the CCA branch is located just superior to the ascending aorta, such that arterial flow in both sides of the CCA is still approximately equivalent to aortic flow velocity prior to bifurcation. The only difference found between between the left and right sides in the tilt study occurred in the A_ĲV_ ^76^. We did not see find significant difference in ĲVP between the left and right sides, hypothesized as due to the fact that both sides sit minimally above CVP.

The main difficulty in detecting significance between the left and right sides is the large intersubject variability. Ogoh *et al.* assessed the difference between the left and right side A_ĲV_ in two conditions: (1) 0° supine with −60 mmHg LBNP; and (2) 60° HUT with no LBNP^44^. This is one of the few studies that assessed both sides of the jugular veins. Similar to our data, Ogoh and colleagues found large variability in their data. With the application of −60 mmHg LBNP, they measured a change in the right A_ĲV_ of −45% ± 49%, and in the left A_ĲV_ of −49% ± 27% (with respect to the supine position and no LBNP). Here, as with our data, the standard deviation of the measurements is too large to draw significant conclusions. This gives us confidence that indeed, in the variables considered, natural variability between subjects is larger than intrasubject differences between the left and right side.

### 4.1 Implications for Countermeasure Design

This study provide new insights into countermeasure design, but it also leads to important questions that warrant future investigation. In particular, related to human spaceflight risks, the three most relevant are: (1) the risk of cardiovascular adaptations contributing to adverse mission performance and health outcomes; (2) the risk of spaceflight-associated neuro-ocular syndrome (SANS); and (3) the concern of venous thromboembolism. Each of these risks are considered briefly in the following paragraphs.

#### 4.1.1 Risk of Cardiovascular Adaptations Contributing to Adverse Mission Performance and Health Outcomes

Since this study only considered acute effects of LBNP, it is difficult to make solid recommendations about a long-term countermeasure for cardiovascular health. We note that the root etiology of the cardiovascular risk is through fluid shifts leading to: (1) alterations in intravascular volume, (2) changes in cardiac and vascular structure/function, (3) oxidative stress, and (4) inflammation^1^. Reducing the headward fluid shift, or at least providing periodic unloading, could prevent or reduce the effect of these causal pathways. The open questions remaining are therefore what are the specific level and protocol of LBNP that are appropriate to mitigate these risks.

This study provides some insight into these questions. First, we note the well-established differences in tolerance between males and females. In our study, we found strong evidence of a difference between the male and female response in absolute variables (e.g., SV, CO) but not in indexed variables (e.g., SI, CI). These results suggest that sex differences are driven principally by differences in anthropometry. This places an upper limit on the level of LBNP that could be used in a spaceflight environment, particularly when the astronauts are in a deconditioned state. From our data, for example, a strength of −50 mmHg caused presyncope in over half of our female subjects. Further analysis may lead to the development of personalized protocols based on anthropometric considerations, rather than a “one size fits all” approach. Second, and most importantly, by considering Figure 2 together with Figure 4, we gain new insight into the relative magnitude of the LBNP *Pressure* effect compared to a gravitationally induced fluid shift (i.e., *Position* effect). Thus, we can use these effect sizes and associated models to determine the appropriate level of LBNP needed to bring back any variable of interest to “Earth normal” conditions when subjected to a headward fluid shift. As one example, we note that a 15° HDT increases cardiac output by 0.35 l/min (89% CrI: 0.26 to 0.46 l/min) with respect to supine levels. In contrast, LBNP reduces CO by 0.046 l/min/mmHg (89% CrI 0.042 to 0.049 l/min/mmHg). Thus, it would appear that only −7.7 mmHg of LBNP are required to return CO to its baseline, supine value. As to be expected, the amount of LBNP required to return to a “baseline” state depends on the variable being considered. For example, for IOP, approximately −29 mmHg of LBNP are required to return IOP to supine levels (when at 15° HDT). Thus, an analysis of which variables are most important to control, combined with further understanding of the long-duration effects, could provide target levels of LBNP for further investigation. It should be noted that there are differences between the physiological response to spaceflight and HDT. In particular, as previously discussed, HDT replicates the fluid shifts but does not remove hydrostatic gradients or alter tissue weight. In addition, the fluid redistribution induced by 15° HDT is most likely of a larger magnitude than the fluid redistribution experienced in true microgravity conditions (which are typically simulated using a 6° HDT paradigm). Thus, countermeasure development will require further investigation and validation of the resultant protocols in microgravity conditions in addition to terrestrial development.

#### 4.1.2 Risk of Spaceflight Associated Neuro-ocular Syndrome (SANS)

In this study, we find that OPP is not influenced by LBNP in the pressure and position ranges measured. A head-down tilt increases OPP, but there is significant evidence for no effect of *Pressure* to subsequently reduce it (%*_ROPE_* = 97.06%). Given the potential relationship between SANS and elevated OPP noted by Petersen *et al.*^57^, and the symptomatic similarities between terrestrial traumatically elevated OPP and SANS^57,106^, LBNP could perhaps not be an effective SANS countermeasure. The potential implications of the effects of LBNP on OPP and, more broadly, SANS, are further discussed in a separate publication by Hall *et al.*^58^. The pathoetiology of SANS is currently unknown, but it is likely the result of multiple contributing factors. Related to fluid pressures, whilst OPP may be important, there are other pressure gradients that also likely play a role, including the translaminar pressure gradient (TLPG)^57^. An important missing piece of information for determining a more complete hemodynamic environment of the ocular system is a measurement of intracranial pressure (ICP). Petersen *et al.* demonstrated that LBNP can reduce ICP in a study using 10 subjects with either parenchymal ICP-sensors or Ommaya-reservoirs fitted to the frontal horn of a lateral ventricle^21^. They found that graded LBNP (in the same range as our experiment) reduced ICP from 15 ± 2 mmHg (0 mmHg LBNP) to 14 ± 4 mmHg (–10 mmHg LBNP), 12 ± 5 mmHg (–20 mmHg LBNP), 11 ± 4 mmHg (–30 mmHg LBNP), 10 ± 3 mmHg (–40 mmHg LBNP), and 9 ± 4 mmHg (–50 mmHg LBNP) (*p <* 0.0001), but that cerebral perfusion pressure (*CPP* = *MAP_mid_*_−_*_brain_* −*ICP*) was unchanged. It is difficult to obtain non-invasive measurements of ICP, but there are a number of existing techniques with varying degrees of accuracy that could be leveraged, including computed tomography and magnetic resonance imaging, transcranial Doppler, electroencephalography power spectrum analysis, and audiological and ophthalmological techniques^107^. Future work should assess the totality of hemodynamic measurements in the head and eyes, including IOP, OPP, ICP, and CPP.

#### 4.1.3 Concern of Venous Thromboembolism

Our results quantify the changes in ĲV blood flow velocity waveform pattern as a function of applied LBNP pressure. In Figure 6 we demonstrated that increasing LBNP can decrease the probability of a Grade 2 or higher flow. Grade 1 and Grade 2 flows are considered normal, whilst the risk of VTE is elevated when attaining Grade 3 and Grade 4 flows^10^. Since this was an acute study, we only observed four instances of Grade 3 Flow in three subjects (two male, one female). In contrast, Marshall-Goebel *et al.* observed Grade 3 or higher flow in 7 of 11 subjects in flight^10^. Thus, our data mainly demonstrate the ability of LBNP to change Grade 2 flow to Grade 1 flow. In order to further understand the suitability of LBNP as a countermeasure to mitigate the concern of VTE, future work should either focus on long duration head-down tilt bed rest (HDTBR) studies and especially spaceflight studies, where Grade 3 or higher flow is more likely to occur.

### 4.2 Discussion of the Bayesian Workflow Methodology

An important contribution of this research effort is the Bayesian workflow used to construct LBNP dose-response curves. We believe that this methodology, which moves away from more common and traditional null hypothesis significance testing (NHST), is better suited to the analysis of both ground-based and spaceflight studies of physiological response, which are often plagued by a lower subject pool. In particular, by removing the reliance on a single value, e.g., *p* ≤ 0.05, to make binary decisions about a null hypothesis, we are able to gather evidence both in favor of, and against, the null and alternative hypotheses. Similarly, whilst many spaceflight studies may find significance, they are often constrained by difficulty in obtaining sufficient power. In a Bayesian framework, power constraints are less important, and it is possible to find evidence even with a low subject pool. Increasing the amount of evidence available reduces the width of the posteriors, increasing our confidence in the estimates, but even a small amount of data are better than no data. Finally, we believe that the Bayesian methodology provides improved understanding of the dose-response parameters. Rather than an often misunderstood interpretation of a confidence interval, the Bayesian credible interval provides estimates as to where the effect sizes lie within the population.

An additional benefit of a Bayesian workflow is that it provides the ability to construct more complicated models that are no longer constrained by the assumptions of (generalized) linear mixed-effects models. Thus, rather than analyzing each variable independently, we constructed a single, highly complex regression model to determine the multivariate structure of the response. The resulting output, in Figure 7, represents a novel understanding of the relationships between the physiological variables considered in the study.

We believe that this Bayesian workflow could be applied to spaceflight studies on physiological response outside of just the cardiovascular system. In particular, it is theoretically possible to elicit the relationship between multiple different physiological systems. For example, this framework will provide insight into questions such as “what is the relationship between cardiovascular degradation and musculoskeletal remodeling during long-duration spaceflight”? Even with the low number of subjects typically included in spaceflight studies, evidence can be built up over time by replacing priors with posteriors from previous studies.

### 4.3 Limitations

There are two key limitations to this study: (1) we only consider the acute response to LBNP, and (2) we are limited to non-invasive measurements. Regarding the acute nature of the study, Lightfoot *et al.* found that adaptation occurs with presyncopal symptom limited LBNP (PSL-LBNP) over the course of a nine-day repeated exposure, with LBNP tolerance increasing 47% over the first five exposures^108^. The authors note an increase in RPP and maximal HR by days 7 and 8, but no change in the MAP response. Lightfoot *et al.* do not comment on adaptation to LBNP when exposed to less than presyncopal strength, but their work notes that adaptation is possible. This indicates that subjects with a low tolerance to LBNP, for example some of our female subjects, may be able to adapt to −50 mmHg LBNP. However, in multiple other studies, LBNP tolerance was found to be highly repeatable in any given individual^19,25,31,109^. When considering LBNP application prior to presyncope, multiple studies by Convertino and Goswami have confirmed that individual cardiovascular response is highly reproducible, even with rest periods as long as one year between tests^17,19,28,30^. This reproducibility also supported our decision to progress, rather than randomize, the presentation of LBNP strength to each subject. In order to determine the utility of LBNP as a spaceflight countermeasure to mitigate SANS or VTE, long term studies must be conducted, either in spaceflight or in HDTBR, to determine whether a periodical unloading of the cephalad fluid shift is able to prevent the manifestation of presyncopal symptoms. Regarding the limitations of non-invasive measurements, a direct measurement of cardiac output could provide the most accurate dose-response relationship as there are observable differences between the results of different methodologies^56,110^. Further, for the autonomic measures, samples of blood plasma catecholamines and other neurohormones along with intracellular magnesium levels^111,112^ could provide further insights into cardiovascular control. Finally, invasive measurement of central venous pressure could provide informative data on cardiac loading conditions and thoracic blood volume^113,114^. The VeinPress device used in the study was chosen for its heritage of use in previous spaceflight and parabolic flight investigations^10,115,116^, thus, facilitating direct comparison between studies. It is acknowledged that invasive measures, such as venous catheterization, may provide more accurate measurements of ĲVP. However, non-invasive measurements using ultrasonography are well-established in a clinical environment^117–119^. The ability to obtain invasive measures could also allow for an accurate measure of ICP^21^. Future studies should attempt to obtain some measure of ICP, which may become easier with advancing technology in the area^107^.

Finally, regarding female subjects, we recorded, but did not control, menstrual phase. Hart *et al.* have noted that sex hormones affect the activity of autonomic pathways on both the sympathetic and parasympathetic sides^120^. Given the fluctuation of hormones across the menstrual cycle, it is envisaged that this could potentially add additional variation into the autonomic response of female subjects, perhaps helping to explain differences in tolerance. Future work should examine the effect of menstrual phase on LBNP tolerability.

## 5 Conclusion

We subjected 24 male and female subjects to graded LBNP to investigate the acute changes in multiple hemodynamic parameters, autonomic indices, and head/neck hemodynamics across the range 0 to −50 mmHg LBNP in both 0° supine and 15° HDT positions. Our data revealed a linear dependence on pressure for all metrics considered except for ĲVP (which presented a logistic dose-response), with varying effect sizes of response. Based on the experimental data collected, we conducted a Bayesian multivariate analysis to construct dose-response curves for all variables across the ranges considered. These dose-response curves demonstrated anthropometrically-driven sex-dependent changes in some metrics related to the systemic hemodynamics, and supported evidence from previous studies regarding different autonomic activation between men and women. For each variable considered, we calculated the relative effect size of a 15° HDT induced cephalad fluid shift, and the LBNP required to counteract it. In particular, we demonstrated the potential for LBNP to reduce jugular venous flow stagnation, and provided a logistic dose-response. Finally, we calculated the relationship structures between all of the variables considered, as well as subject characteristics, finding correlation structures between many groups of variables. These findings provide data to support spaceflight countermeasure development to mitigate the risk of cardiovascular degradation, venous thromboembolism events, and SANS (although the lack of effect of LBNP on OPP warrants further investigation).

### Appendix A Raw Data

#### A.1 Systemic Hemodynamic Response

Figure A1 shows the evolution of systemic hemodynamic parameters (mean ± SE) as a function of LBNP pressure (the seated baseline has been removed for clarity). All measured variables follow an approximately linear trend with respect to LBNP pressure.

**Figure A1:**
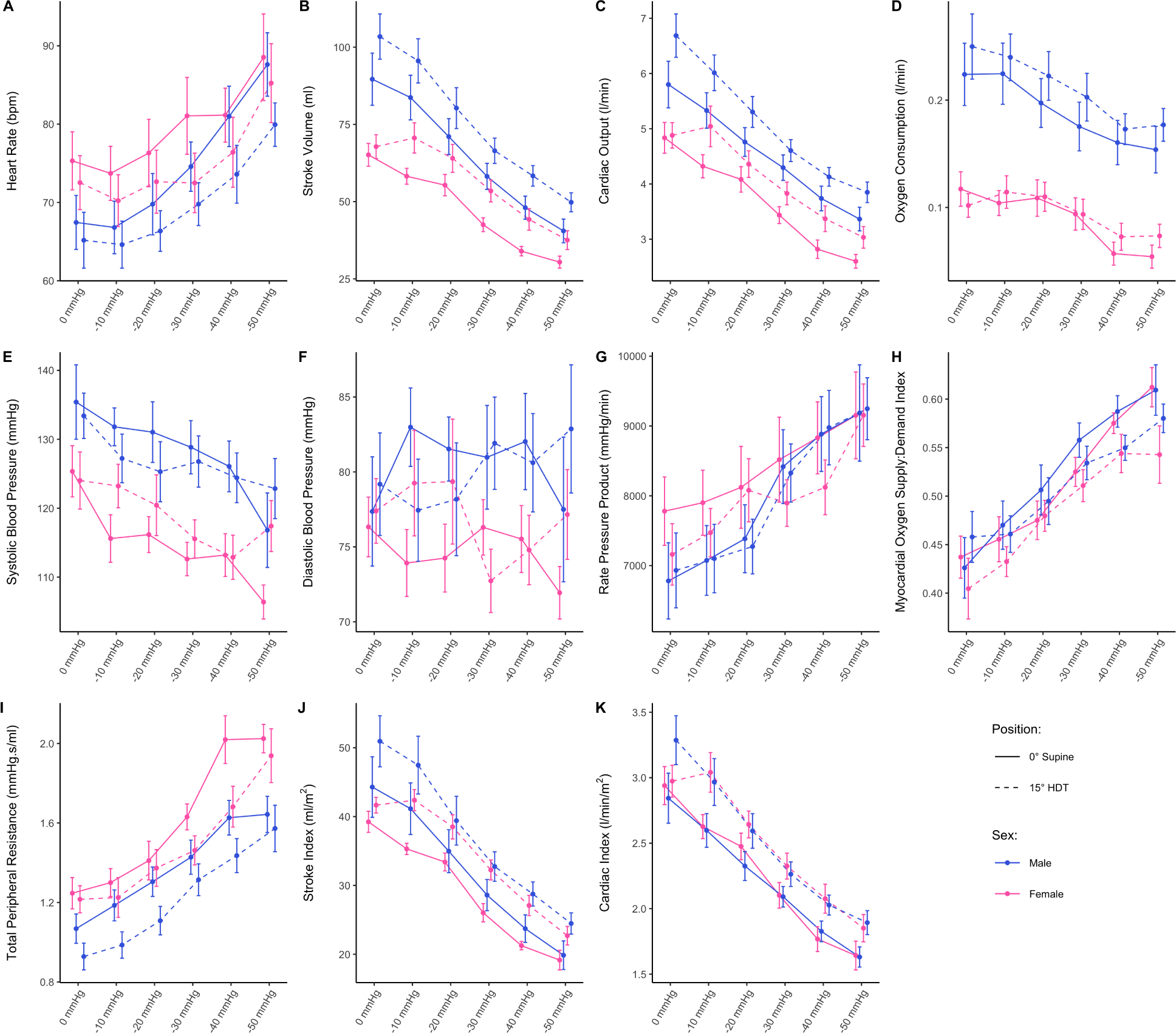
**(A-K)** Systemic hemodynamic variables as a function of LBNP pressure in 0° supine (solid line) and 15° HDT (dashed line) positions, collected on 24 subjects (12 male, 12 female). Measurements were taken at 0 mmHg, −10 mmHg, −20 mmHg, −30 mmHg, −40 mmHg, and −50 mmHg of LBNP. Data are presented as means ± SE at each pressure level. **(A)** HR, heart rate; **(B)** SV, stroke volume; **(C)** CO, cardiac output; **(D)** VO2, oxygen consumption; **(E)** SBP, systolic blood pressure; **(F)** DBP, diastolic blood pressure; **(G)** RPP, rate pressure product; **(H)** MO, myocardial oxygen supply:demand index; **(I)** TPR, total peripheral resistance; **(J)** SI, stroke index; **(K)** CI, cardiac index.

In males, heart rate (HR, Figure A1A) increases from 67.5 ± 3.4 bpm at 0 mmHg of LBNP to 87.6 ± 4.1 bpm at −50 mmHg of LBNP in 0° supine, and from 65.2 ± 3.6 bpm at 0 mmHg of LBNP to 79.9 ±2.8 bpm at −50 mmHg of LBNP in 15° head down tilt (HDT). In general, female subjects have a HR that is 2.9 bpm (89% CrI: −3.4 to 10.1 bpm) higher than males (*pd* = 77.83%, %*_ROPE_* = 19.41%). Stroke volume (SV, Figure A1B) and cardiac output (CO, Figure A1C) decrease at an average rate (males and females together) of 8.7 ml (89% CrI: 8.2 to 9.4 ml) and 0.46 l/min (89% CrI: 0.42 to 0.49 l/min), respectively, for every 10 mmHg increase in LBNP strength (more negative). In absolute values, SV and CO are higher in males, with respect to the females, by 24.9 ml (89% CrI: 16.9 to 33.6 ml) and 0.62 l/min (89% CrI: 0.21 to 1.06 l/min), respectively. However, when variables are indexed by body surface area (BSA), cardiac index (CI, Figure A1K) is equivalent in males and females, decreasing by 0.26 l/min/m^2^ (89% CrI: 0.24 to 0.28 l/min/m^2^, males and females together) per 10 mmHg increase in LBNP. On the other hand, stroke index (SI, Figure A1J) is still higher in male subjects by, on average, 4.3 ml/m^2^ (89% CrI: 0.4 to 8.4 ml/m^2^).

Systolic blood pressure (SBP, Figure A1E), which is higher in males by 7.4 mmHg (89% CrI: 0.9 to 13.8 mmHg), decreases slightly at an average rate of 1.9 mmHg (89% CrI: 1.3 to 2.6 mmHg) per 10 mmHg increase in LBNP. However, diastolic blood pressure (DBP, Figure A1F) appears to hold a relatively constant value with no clear trend.

Rate pressure product (RPP, Figure A1G), myocardial oxygen supply:demand index (MO, Figure A1H), and total peripheral resistance (TPR, Figure A1I) increase linearly across the range of LBNP values considered. There is no difference in MO between male and female subjects (0.01, 89% CrI: −0.04 to 0.05); however, RPP and TPR appear to be slightly higher in females (for RPP: 270 mmHg/min higher, 89% CrI −600 to 1200 mmHg/min, *pd* = 70.06%, %*_ROPE_* = 22.47%; for TPR: 0.15 mmHg.s/ml higher, 89% CrI −0.01 to 0.30 mmHg.s/ml, *pd* = 93.69%, %*_ROPE_* = 10.02%). In RPP, this is most noticeable at lower pressure levels, and is likely driven by the higher resting HR in female subjects.

The largest difference between males and females appears in oxygen consumption (VO2), where males consume twice as much oxygen as the females. Males have an average VO2 of 0.22 ± 0.03 l/min at 0 mmHg of LBNP, and this value decreases to 0.15 ± 0.02 l/min at −50 mmHg of LBNP (in 0° supine). Females have an average VO2 of 0.12 0.02 l/min at 0 mmHg of LBNP, and this value decreases to 0.05 ± 0.01 l/min at −50 mmHg of LBNP (0° supine). Overall, the effect of LBNP, sex, or position (0° supine or 15° HDT) appear minimal. This result is supported by evidence from our previous tilt studies^77^, which noted no effect of tilt angle on oxygen consumption.

#### A.2 Autonomic Response

Figure A2 shows the evolution of autonomic parameters (mean ± SE) as a function of LBNP pressure (the seated baseline has been removed for clarity).

**Figure A2:**
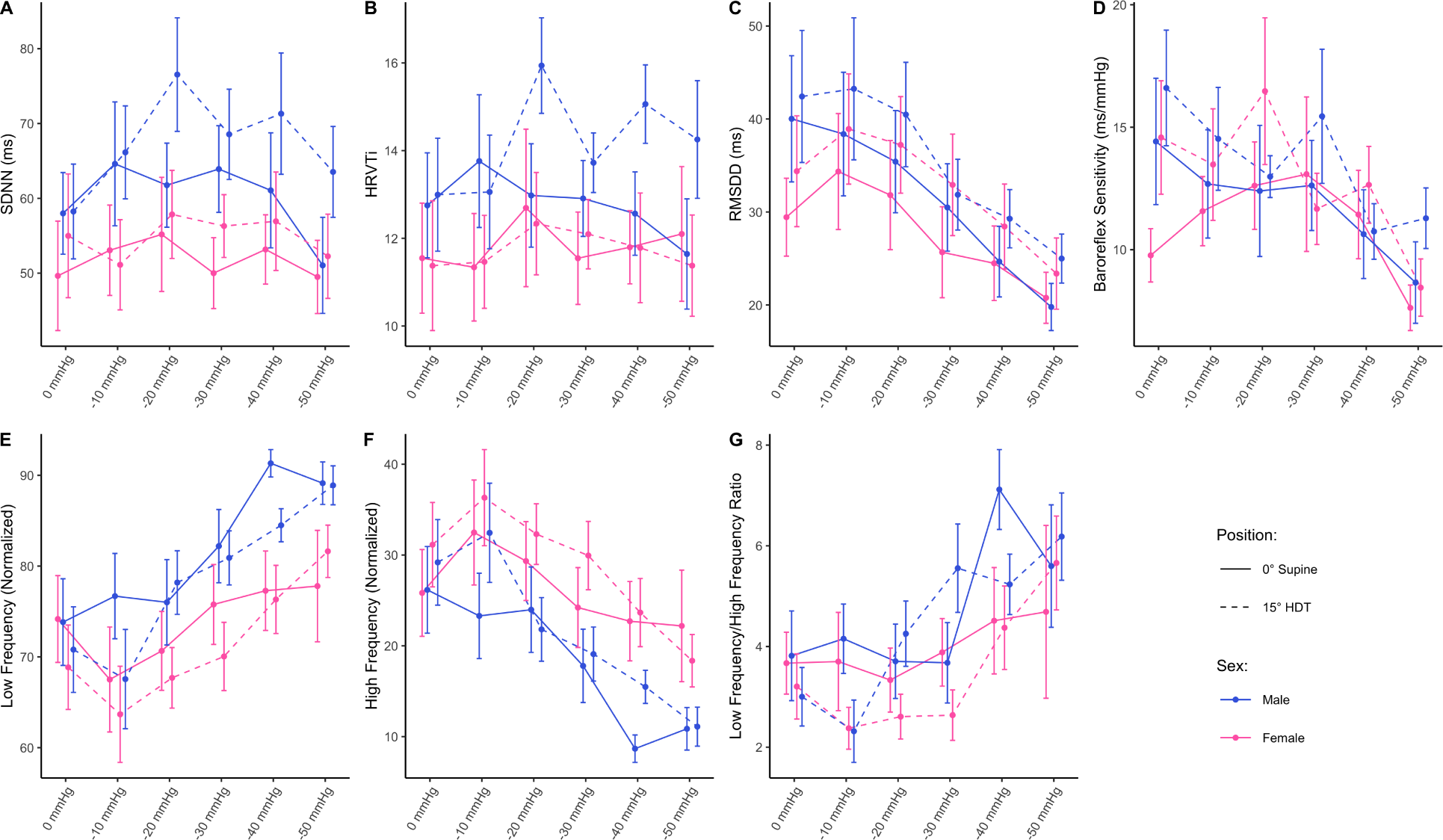
**(A-G)** Autonomic variables as a function of LBNP pressure in 0° supine (solid line) and 15° HDT (dashed line) positions, collected on 24 subjects (12 male, 12 female). Measurements were taken at 0 mmHg, −10 mmHg, −20 mmHg, −30 mmHg, −40 mmHg, and −50 mmHg of LBNP. Data are presented as means ± SE at each pressure level. **(A)** SDNN, standard deviation of NN intervals; **(B)** HRVTi, heart rate variability triangular index; **(C)** RMSDD, root mean square of direct differences of NN intervals; **(D)** BRS, baroreflex sensitivity; **(E)** LFNorm, normalized low frequency power spectral density; **(F)** HFNorm, normalized high frequency power spectral density; **(G)** LF/HF, low frequency to high frequency ratio.

Broadly, there is minimal effect of LBNP on overall heart rate variability, as evidenced by minimal significant effect of LBNP on SDNN (Figure A2A) or HRVTi (Figure A2B). On the other hand, the overall balance of sympathetic and vagal activity is clearly altered by LBNP. In particular, RMSDD (Figure A2C), a marker of vagal activity, decreases from 35.5 ± 3.0 ms at 0 mmHg of LBNP to 22.4 ± 1.5 ms at −50 mmHg of LBNP (average of both sexes and both positions), a reduction of 2.8 ms (89% CrI: 2.1 to 3.5 ms) per 10 mmHg increase in LBNP. This decrease in vagal activity is further supported by the reduction in normalized high frequency power spectral density (Figure A2F) from 28.1 ± 2.3 at 0 mmHg of LBNP to 14.5 ± 1.6 at −50 mmHg of LBNP (average of both sexes and both positions). In addition, this reduction in vagal activity is further matched by a corresponding increase in sympathetic activity supported by the increase in normalized low frequency power spectral density (Figure A2E) and the increase in the low/high frequency ratio (Figure A2G), which is a marker of sympathovagal balance.

Finally, baroreflex sensitivity (Figure A2D) decreases slightly with LBNP, from 13.8 ± 1.1 ms/mmHg at 0 mmHg to 9.2 ± 0.7 ms/mmHg at −50 mmHg (average of both sexes and both positions). This is an average decrease of 0.9 ms/mmHg (89% CrI: 0.6 to 1.2 ms/mmHg) per 10 mmHg increase in LBNP. This reduction in sensitivity is likely related to the reduction in blood flow and pressure in the carotid sinus and aortic arch with increased LBNP, as blood is pooled in the lower body.

#### A.3 Head/Neck Response

Figure A3 shows the evolution of head/neck parameters (mean ± SE), excluding ĲVF, as a function of LBNP pressure. Figure A4 shows the ĲV blood flow velocity waveform pattern as a function of LBNP pressure. The seated baseline data have been removed for clarity.

**Figure A3:**
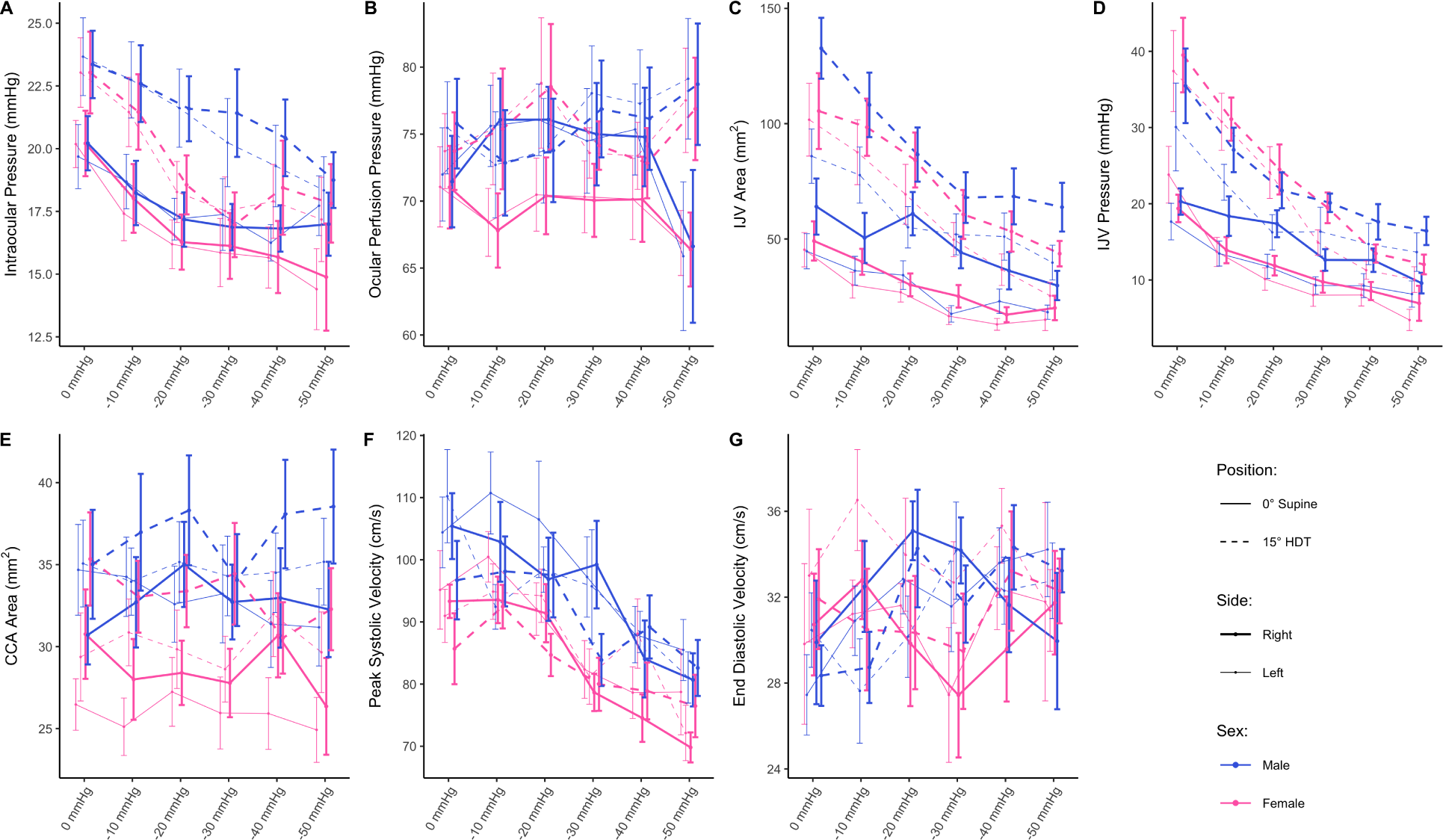
**(A-G)** Head/neck variables as a function of LBNP pressure in 0° supine (solid line) and 15° HDT (dashed line) positions, collected on 24 subjects (12 male, 12 female). Thick lines represent the right side and thin lines represent the left side. Measurements were taken at 0 mmHg, −10 mmHg, −20 mmHg, −30 mmHg, −40 mmHg, and −50 mmHg of LBNP pressure. Data are presented as means ± SE at each pressure level. **(A)** IOP, intraocular pressure; **(B)** OPP, ocular perfusion pressure; **(C)** A_ĲV_, internal jugular vein cross sectional area; **(D)** ĲVP, internal jugular vein pressure; **(E)** A_CCA_, common carotid artery area; **(F)** PSV, peak systolic velocity; **(G)** EDV, end diastolic velocity.

**Figure A4:**
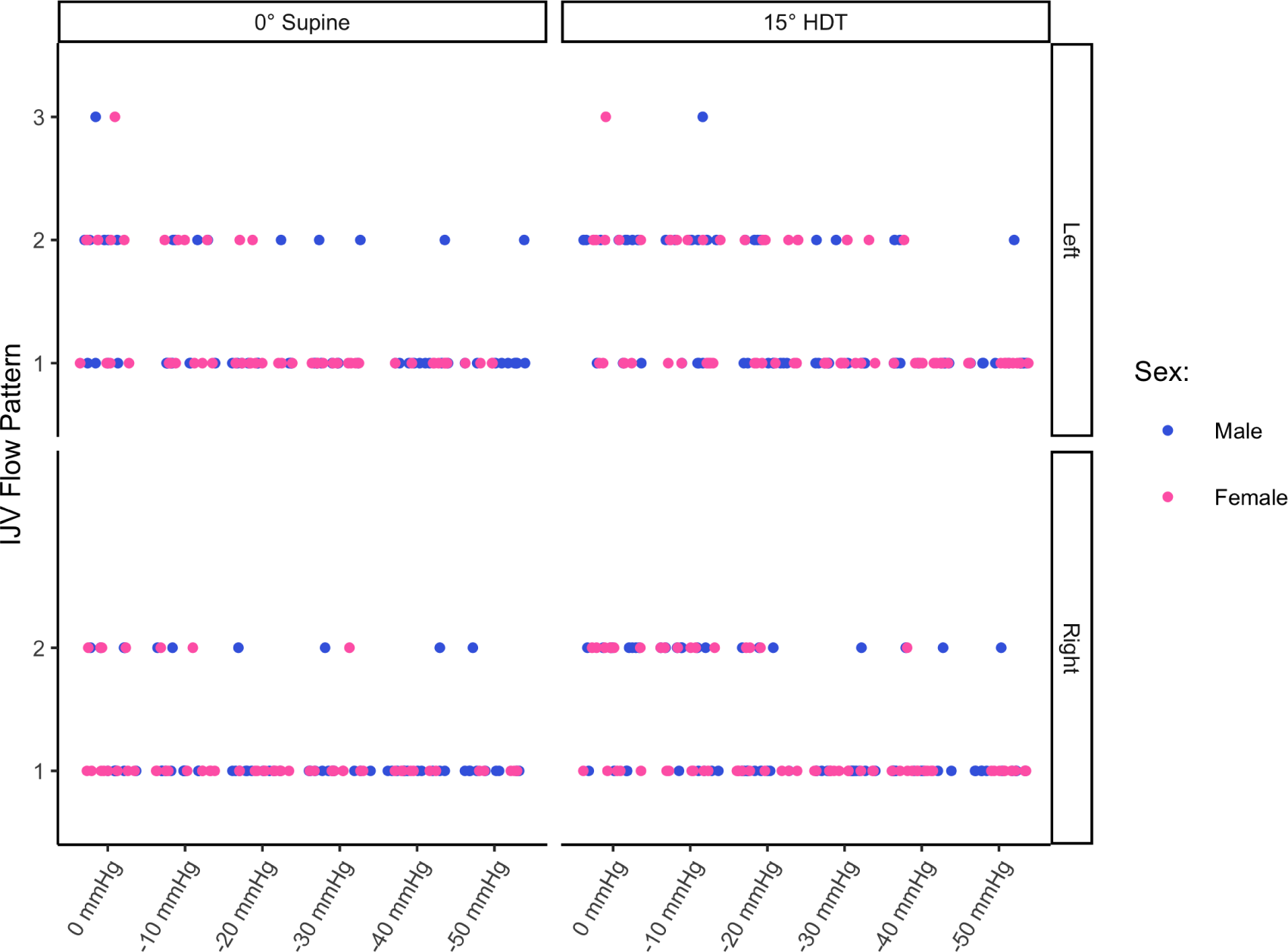
Internal jugular vein (ĲV) blood flow velocity waveform pattern as a function of LBNP pressure in 0° supine (left column) and 15° HDT positions (right column), on the left (top row) and right (bottom row) sides. Data were collected on 24 subjects (12 male, 12 female). Flow grade patterns are described in the main text, illustrated in Figure 1, and taken from Marshall-Goebel *et al.*^10^.

There appears to be little difference between the right and left sides (thick and thin lines, respectively) in any of the variables considered. In the 15° HDT position, intraocular pressure (IOP, Figure A3A) is 2.9 mmHg (89% CrI: 2.3 to 3.4 mmHg) higher than IOP in the 0° supine position. IOP also decreases linearly from 20.1 ± 0.6 mmHg at 0 mmHg of LBNP to 16.4 ± 0.7 mmHg at −50 mmHg of LBNP (0° supine, average of both sexes and both sides). In 15° HDT, IOP decreases from 23.3 ± 0.7 mmHg at 0 mmHg of LBNP to 18.1 ± 0.7 mmHg at −50 mmHg of LBNP (both sexes and sides averaged). In contrast, there appears to be no significant effect of LBNP on ocular perfusion pressure (OPP, Figure A3B), with the response remaining relatively constant across all LBNP levels considered (there is even a slight increase in 15° HDT from 74.7 ± 1.5 mmHg at 0 mmHg of LBNP to 78.1 ± 2.1 mmHg at −50 mmHg of LBNP (male and female, both sides)).

The common carotid artery area (CCA, Figure A3E) does not show a significant effect with LBNP, although we find a small effect of position. The average A_CCA_ at 0° supine is 30.3 ±0.5 mm^2^, which increases to 33.7 ±0.6 mm^2^ in 15° HDT (average of all pressure levels, sexes, and sides). There is minimal change in end-diastolic velocity of the CCA with LBNP (Figure A3G); however, the peak systolic velocity (Figure A3F) decreases from 97.7 ± 2.1 cm/s at 0 mmHg of LBNP to 79.1 ± 1.7 cm/s at −50 mmHg of LBNP (average of both side, both sexes, and both positions), indicating an overall decrease in the pulse velocity with LBNP strength (more negative).

Finally, the internal jugular vein (ĲV) exhibits a strong response to LBNP, with decreases of 8.8 mm^2^ (89% CrI: 7.3 to 10.2 mm^2^) per 10 mmHg of LBNP in area (A_ĲV_, Figure A3C) and 2.7 mmHg (89% CrI: 2.4 to 3.1 mmHg) per 10 mmHg of LBNP in pressure (ĲVP, Figure A3D). Both A_ĲV_ and ĲVP increase in the 15° HDT position with respect to the 0° supine position: A_ĲV_ increases by 40.1 mm^2^ (89% CrI: 35.5 to 44.8 mm^2^) and ĲVP increases by 7.9 mmHg (89% CrI: 6.9 to 9.0 mmHg). However, results do not show an effect of side on A_ĲV_. These results contrast to our previous data from tilt studies^76^. However, this is not necessarily surprising, since the larger differences in A_ĲV_ between the left and right sides only begin to appear in HDT and continue to get amplified at larger tilt angles (30° HDT and 45° HDT). In the present study, subjects remained at 15° HDT during the entire intervention, which can explain the lack of significant differences in A_ĲV_between the right and left sides. With respect to the ĲV blood flow velocity waveform, Figure A4 reveals that, whilst the majority of observed flows are at Grade 1, we find more instances of Grade 2 flow at lower LBNP levels (and even two cases of Grade 3 flow stagnation, both appearing in the left ĲV). Overall, our results suggest that LBNP is effective at reducing the instances of Grade 2 flow.

## Acknowledgements

The authors would like to thank our anonymous subjects for their willingness to participate in the study and the members of the Texas A&M Bioastronautics and Human Performance Laboratory for their assistance. In addition, we would like to thank Technavance Inc., especially Nathan Harrison, for all their technical support with the Lower Body Negative Pressure (LBNP) chamber.

## Availability of data and material

The datasets analyzed for this study are publicly available, a repository can be found on GitHub: https://github.com/BHP-Lab/LBNP-Dose-Response

## Code availability

Not applicable.

## Authors’ contributions

RSW and AD-A contributed to the study conception and design. Material preparation, data collection, and analysis were performed by RSW and AD-A. The first draft of the manuscript was written by RSW and commented on by AD-A. All authors read and approved the final manuscript.

## Ethics approval

All procedures performed in studies involving human participants were in accordance with the ethical standards of the institutional and/or national research committee and with the 1964 Helsinki Declaration and its later amendments or comparable ethical standards. The study was approved by the Human Research Protection Program of Texas A&M University with Institutional Review Board number IRB2020-0724F.

## Consent to participate

Informed consent was obtained from all individual participants included in the study.

## Consent for publication

Not applicable.

## Footnotes

i Kruschke suggests ±0.1 as a default value for a standardized parameter ^48^, equivalent to a negligible effect size according to Cohen ^49^.

ii Compared to correlation in a frequentist framework, the maximum a-posteriori estimate is analogous to the *r*-value, whilst the significance, i.e., whether the posterior distribution encompasses 0, is analogous to the *p*-value.

